# Complex I Inhibition Redirects Autophagy from Bulk Turnover to Mitochondrial Clearance

**DOI:** 10.64898/2026.06.04.730191

**Authors:** Thuraya M. Mutawi, Shweta R. Malvankar, Andrea Graziani, Udita Shah, Khanh Nguyen, David G. Broadbent, Zachary Clark, Michaella J. Rekowski, Susan M. Lunte, Carlo Barnaba

**Affiliations:** Department of Pharmaceutical Chemistry, University of Kansas, Lawrence, KS, USA; Department of Molecular Biosciences, University of Kansas, Lawrence, KS, USA; School of Pharmacy, University of Kansas, Lawrence, KS, USA; College of Osteopathic Medicine, Michigan State University, East Lansing, MI, USA; Mass Spectrometry/Proteomics Core Laboratory, University of Kansas Medical Center, Kansas City, KS, USA; Department of Cancer Biology, University of Kansas Medical Center, Kansas City, KS, USA; Department of Chemistry, University of Kansas, Lawrence, KS, USA; Ralph N. Adams Institute for Bioanalytical Chemistry, University of Kansas, Lawrence, KS, USA

**Keywords:** autophagy initiation, mitophagy, mitochondrial complex I, ULK1 kinase, metformin

## Abstract

Mitochondria are both a major autophagic substrate and a principal source of signals that control autophagy, yet it is unknown whether a mitochondrial signal can specify which substrate the pathway degrades. Using metformin to inhibit Complex I in human cells, we find that autophagy is redirected rather than induced or suppressed. Inhibition shifts the NAD^+^/NADH ratio before perturbing the adenylate pool, separating the redox arm of the lesion from the bioenergetic arm. Single-molecule tracking of endogenous ATG13 and WIPI2 shows that initiation is preserved while WIPI2 tethering fails, generating shortlived, abortive autophagosomes; the ULK1 interactome concurrently disengages from cytoskeletal scaffolds, and Parkinindependent mitolysosomes accumulate. This response differs from mTORC1 inhibition and does not require PEN2, and AMPK is engaged non-canonically: α1/α2 restrain ATG13 at initiation, while AMPK activation alone does not reproduce the WIPI2 phenotype. Occurring at therapeutic concentrations, these results identify the ULK1 complex as the step at which Complex I inhibition reassigns autophagic substrate.

## Introduction

Autophagy is a catabolic pathway that delivers cytosolic material to the lysosome for degradation. Autophagy can be non-selective, capturing cytoplasm in bulk, or selective, targeting defined substrates such as damaged organelles and protein aggregates. ^1,2^ Both forms use the same core machinery but differ in what they degrade, and the physiological consequences of autophagy depend on which predominates. In cancer cells, elevated basal autophagy supports survival under nutrient limitation, and researchers have pursued pharmacological modulation of the pathway as a therapeutic strategy. ^3^ Autophagosome formation begins when the ULK1 kinase complex – ULK1, ATG13, FIP200/RB1CC1, and ATG101 – is recruited to nascent assembly sites. ^4^ ULK1 activation triggers a phospholipid signaling cascade that generates PI3P on ATG9-containing vesicles, which serve as the seed of the phagophore. ^4^ PI3P is bound by WIPI2, which recruits the ATG16L1-ATG12-ATG5 complex and allows conjugation of ATG8-family proteins (LC3, GABARAP) to the growing membrane. Initiation and maturation are therefore separable steps, and a phagophore can initiate without being tethered; in that case, it fails to expand and is shortlived. ^4^ Bulk measurements of LC3-II do not distinguish these outcomes, because LC3-II accumulates whether autophagosomes are produced more rapidly or cleared more slowly.

How this machinery responds to a mitochondrial metabolic signal is less well defined. Metformin inhibits mitochondrial Complex I, an activity characterized in isolated mitochondria and intact cells. ^5–7^ Complex I inhibition has two consequences. It prevents NADH reoxidation and raises the cellular NADH/NAD^+^ ratio, and it restricts ubiquinone-coupled electron flux and, with it, oxidative ATP synthesis. These are usually treated together as energy stress, but they are distinct signals and need not act on the same timescale. Which of them a given cellular response depends on remains unclear. The route by which metformin reaches AMPK has also been debated. AMPK senses the AMP/ATP ratio and activates catabolic pathways while suppressing biosynthetic ones. In the classical model, Complex I inhibition depletes cellular ATP and activates AMPK through cytoplasmic AMP sensing by the AXIN-LKB1 complex. ^5–8^ A parallel route was proposed in which the γ-secretase subunit PEN2 bridges v-ATPase inhibition to AXIN/LKB1 recruitment at the lysosomal surface, enabling AMPK activation at clinically relevant (5–50 µM) metformin concentrations independently of cellular AMP. ^8,9^ The two models make opposite predictions for autophagy. Complex I inhibition drives broad AMP-AMPK signaling but spares the lysosome; the PEN2 model centers on v-ATPase inhibition – an outcome that should impair the very lysosomal degradation autophagy depends on. In addition, NAD^+^ has recently been identified as a direct allosteric regulator of AMPK at the γ1-CBS1 site, capable of activating the kinase in the absence of changes in AMP/ATP, ^10^ which would allow the redox arm of Complex I inhibition to signal on its own. Distinguishing between these routes has remained difficult without cellular systems that resolve metformin’s action at molecular resolution.

The dominant paradigm holds that AMPK activation inhibits mTOR to stimulate autophagy, with reported induction across several cellular models. ^11–13^ A converging body of recent work has begun to invert this interpretation. Park et al. demonstrated that AMPK activation under energy stress drives ULK1 inactivation and net suppression of autophagic activity; ^14^ Kazyken et al. independently reported AMPK-mediated autophagy suppression during prolonged amino acid deprivation, ^15^ and we mapped the downstream lesion to the phagophore-to-autophagosome maturation step, showing that AMPK impairs WIPI2 tethering at the nascent membrane and generates short-lived, abortive autophagosomes. ^16^ Previous work, however, left open whether the same tethering lesion is reached by perturbing mitochondrial metabolism rather than by withdrawing a nutrient, whether a specific AMPK subunit carries the signal, and – most consequentially – what the initiation machinery engages instead once bulk tethering fails. We take up these questions here, combining targeted metabolomics, immunoprecipitation-mass spectrometry of endogenously HaloTagged ULK1, and singleparticle tracking of core autophagy factors in living human osteosarcoma cells. Metformin retains Complex I inhibitory activity at therapeutic concentrations, progressively depleting the cellular nucleotide pool. Within hours, the ULK1 complex undergoes a non-canonical reprogramming that selectively displaces WIPI2 from membrane-tethered states and generates short-lived, abortive autophagosomes. Genetic dissection shows that this response operates independently of the PEN2-lysosomal route ^9^, the AMPK-activating pathway recently proposed as metformin’s primary cellular target, and that AMPK is engaged in a subunit-specific manner that separates autophagy initiation and maturation. Bulk autophagy is suppressed in parallel with engagement of selective mitochondrial clearance. The autophagy machinery is therefore not switched off but redirected from cytosolic substrate turnover toward mitochondria through reprogramming of the ULK1 complex, distinct from the route metformin is thought to engage in its lysosomal mode of action. Because autophagy and mitochondrial quality control are now central to metformin’s long-term metabolic effects in type 2 diabetes and to its emerging activity against tumor cells in which basal autophagy supports survival, ^17,18^ identifying ULK1 as that node has direct implications for both indications and for the rational design of pharmacological therapies.

## Results

### Metformin-mediated Complex I inhibition precedes adenylate-pool collapse

Whether metformin’s Complex I inhibitory activity, originally characterized at supratherapeutic concentrations, ^7^ is engaged at clinically relevant doses now centered in the PEN2 model ^9^ remains an open question, and one whose answer constrains every downstream model of metformin signaling. To define the metabolic consequences of metformin exposure, we quantified 11 intracellular metabolites at 2, 6, and 24 h in U2OS cells treated with metformin (5, 200, or 500 µM) or with the mitochondrial uncoupler CCCP (1 µM) as a positive control for bioenergetic disruption (Fig. 1a–f). The earliest detectable response was redox, not energetic. At 2 h, ATP and ADP levels were preserved at clinically relevant doses (Met 5 and Met 200 µM; n.s. vs CTRL), yet NADH had already doubled in the Met 500 group (1.20 ± 0.08 to 2.64 ± 0.39 nmol/mg, p < 0.001) and NAD^+^ had begun to decline (9.90 ± 0.48 vs 12.68 ± 0.50, p < 0.001; Fig. 1d). Even at Met 5 µM, the NAD^+^/NADH ratio was already significantly reduced at 2 h (8.78 ± 0.59 vs 10.60 ± 0.92 in CTRL; p < 0.05), in the absence of any change in adenylate species. The NAD^+^/NADH ratio is therefore the most sensitive readout of metformin’s mitochondrial action, registering Complex I inhibition before the adenylate pool is perturbed. By 24 h, the ratio collapsed to *∼*1, comparable to CCCP. Energy-pool perturbations followed a delayed and dose-dependent trajectory. At 6 h, the Met 500 group exhibited concomitant declines in ATP (29.56 ± 1.45) and ADP (11.79 ± 0.67), marking the limit of adenylate-kinase compensation. By 24 h, even clinical-dose metformin (5 µM) produced substantial adenylate-pool contraction (ATP 31.87 ± 0.86 vs CTRL 55.00 ± 1.62; ADP 7.83 ± 0.34 vs CTRL 30.03 ± 0.62; both p < 0.001), a finding that is at variance with prior reports of stable adenylate species at this dose in primary mouse hepatocytes ^9^ and that we attribute to the higher mitochondrial reserve capacity of hepatocytes relative to U2OS osteosarcoma cells. ^19^ Inosine, the terminal product of purine catabolism, emerged as the most informative marker of this collapse, rising at 6 h (Met 500: 4.93 ± 0.42; CCCP: 9.33 ± 0.86 nmol/mg; both p < 0.001) and reaching 15.83 ± 0.65 (Met 500) and 22.00 ± 1.20 (CCCP) by 24 h. The inosine response was monotonic across the dose range and detectable even at 5 µM (4.4-fold over control, p < 0.001), establishing nucleotide catabolism as a sensitive index of metformin exposure at therapeutic concentrations.

**Fig. 1.**
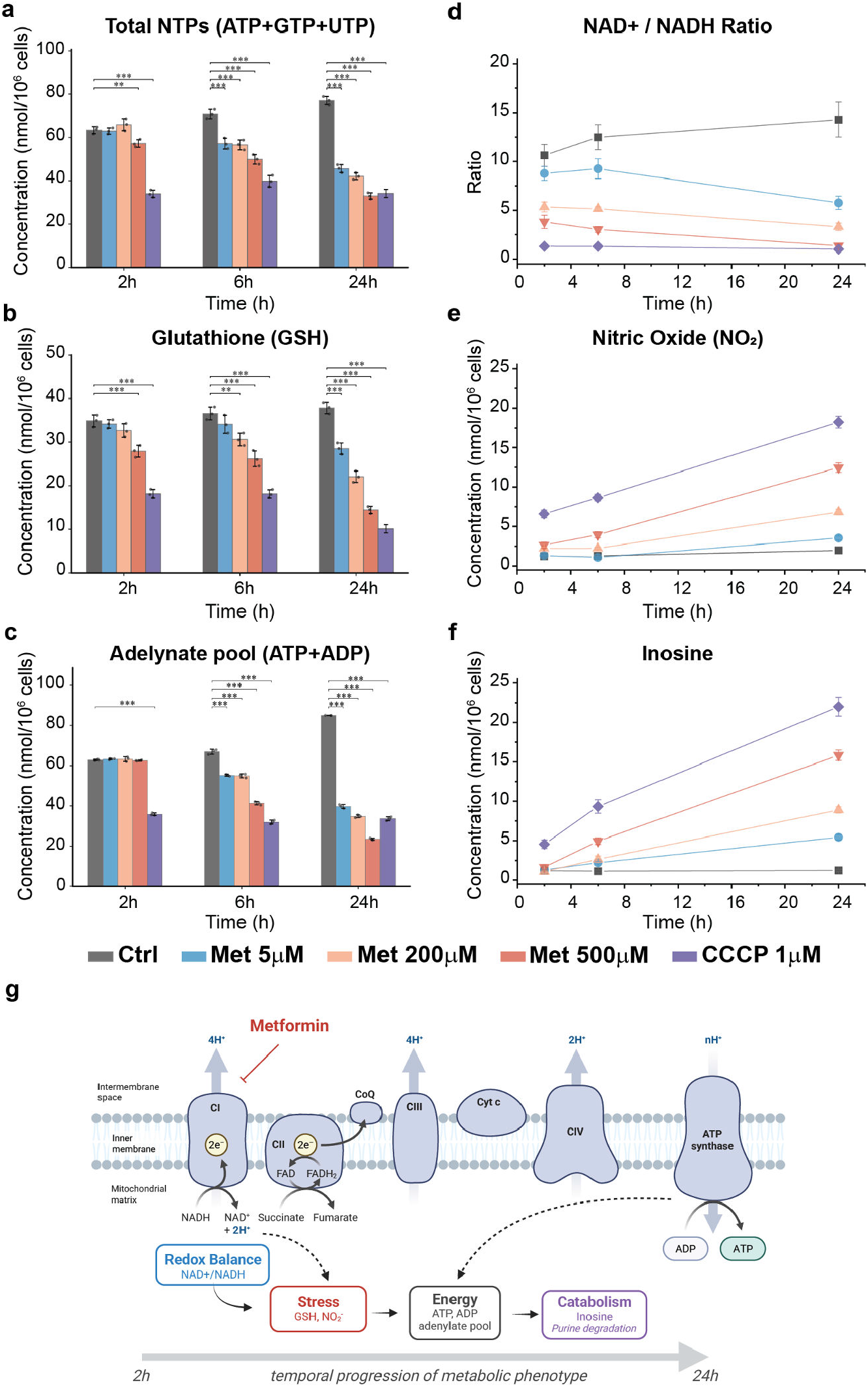
Complex I inhibition by metformin precedes adenylate-pool collapse. **a–f**, LC-based quantification of intracellular metabolites in U2OS cells treated with vehicle (Ctrl), metformin (5, 200, or 500 µM), or the mitochondrial uncoupler CCCP (1 µM) at 2, 6, and 24 h. **a**, Total nucleoside triphosphate pool (ATP + GTP + UTP). **b**, Reduced glutathione (GSH). **c**, Adenylate pool (ATP + ADP). **d**, NAD^+^/NADH ratio. **e**, Nitrite (NO_2_^−^) as a surrogate for nitric oxide production. **f**, Inosine, the terminal product of purine catabolism. **g**, Schematic summarizing the temporal progression of metabolic responses to metformin: Complex I inhibition perturbs the NAD^+^/NADH redox balance first (2 h), followed by oxidative-nitrosative stress (GSH depletion, nitrite accumulation; 6–24 h), then energy-pool failure (adenylate decline; 6–24 h), and finally nucleotide catabolism (inosine accumulation; 6–24 h). Created with Biorender.com. Bar graphs in **a-c** and **i-l** show mean ± 1SD. Three biological replicates per condition. Metabolite concentrations are normalized to cell number and expressed as nmol per million cells. Statistical comparisons versus Ctrl within each time point in **a-c** are by one-way ANOVA with Dunnett’s post-hoc test; \**p* < 0.05, \*\**p* < 0.01, \*\*\**p* < 0.001.

Sustained mitochondrial impairment translated into oxidative and nitrosative stress, with kinetics that distinguished clinical from supratherapeutic doses. At 2 h, glutathione (GSH) was preserved at clinical doses (Met 5: 34.18 ± 0.98; Met 200: 32.67 ± 1.56 vs CTRL 34.88 ± 1.38; n.s.) but already significantly depleted at Met 500 (27.90 ± 1.35; p < 0.001) and severely compromised under CCCP (18.17 ± 0.96; p < 0.001), indicating that high-dose metformin and acute uncoupling rapidly outpace thiol-defense capacity. Clinical-dose GSH depletion emerged at later time points (**Fig. 1b**). Nitrite accumulated more slowly, reaching 12.47 ± 0.60 in Met 500 by 24 h. The coincidence of GSH collapse and nitrite accumulation at 24 h is consistent with progressive oxidative-nitrosative stress downstream of sustained mitochondrial dysfunction. Across all four metabolic axes (adenylate pool, NAD redox, thiol defense, and nucleotide catabolism), the Met500 trajectory at 24 h closely paralleled that of acute CCCP-induced uncoupling, supporting a primary mechanism of metformin action through mitochondrial impairment, with effects detectable at clinically relevant doses on the most sensitive readouts.

### Complex I inhibition uncouples productive autophagic flux from bulk LC3 accumulation through a selective autophagosome tethering defect

Static LC3-II Western blots cannot distinguish enhanced autophagosome biogenesis from impaired lysosomal clearance and have produced contradictory reports of autophagy induction and suppression. To resolve Complex I inhibition of autophagy at molecular resolution, we used an endogenous HaloTag reporter system ^4^ to image autophagosome biogenesis in U2OS cells treated with metformin, followed by a multimodal foci analysis pipeline (K-FOCUS). ^16^ Two cell lines were tested, in which either ATG13 or WIPI2 was tagged with HaloTag at its native locus and stably expressed alongside GFP-LC3. This dual-channel system reports three sequential stages of autophagosome assembly: initiation (ATG13), membrane tethering (WIPI2), and lipidation (LC3) at endogenous protein stoichiometry, with each focus scored as a single, time-resolved event (**Fig. 2a, c; Movies S1-8**). Cells were treated across a 100-fold metformin dose range (5, 200, 500 µM) that spans sub-therapeutic to supraphysiological concentrations, ^9^ in combination with bafilomycin A1 (Baf) to block lysosomal turnover and the direct AMPK activator O-304^20^ to test whether AMPK activation alone reproduces or rescues metformin’s effect.

**Fig. 2.**
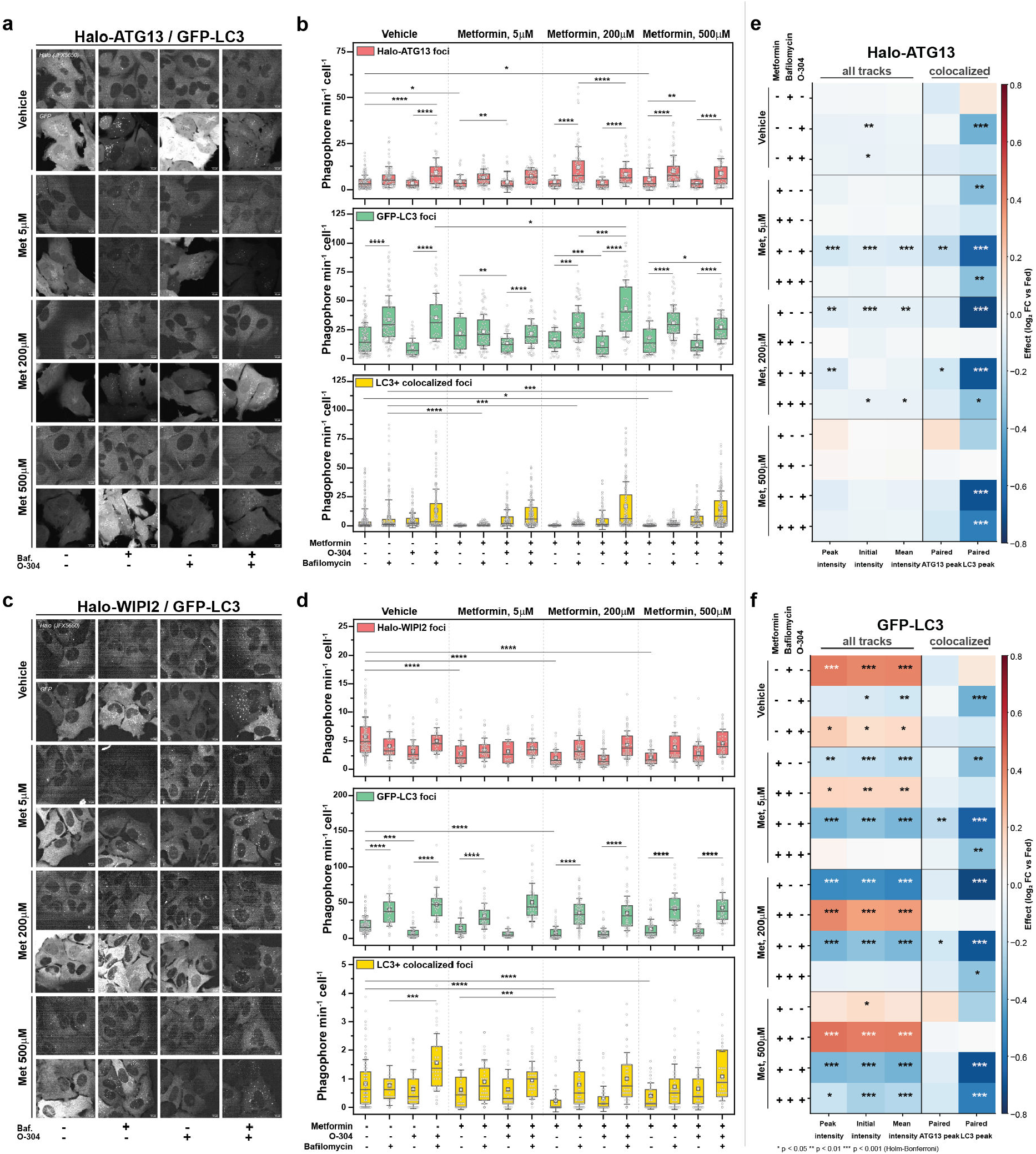
Metformin selectively suppresses WIPI2-tethered, productive autophagosome assembly while sparing ATG13 initiation. **a**, Representative live-cell images of U2OS cells co-expressing endogenously HaloTagged ATG13 (JFX650, top) and stably integrated GFP-LC3 (bottom), treated with vehicle or metformin (5, 200, or 500 µM) for 2 h, in combination with the direct AMPK activator O-304 (1 µM) and the v-ATPase inhibitor bafilomycin A1 (Baf, 100 nM) as indicated. Scale bar, 10 µm. **b**, Per-cell quantification of autophagic foci in the Halo-ATG13 / GFP-LC3 line: Halo-ATG13 initiation foci (top, red), GFP-LC3 lipidated foci (middle, green), and ATG13^+^/LC3^+^ colocalized foci reporting on productive autophagosome assembly (bottom, yellow). **c**, Representative live-cell images for the parallel reporter line, in which WIPI2 was endogenously HaloTagged at its native locus and co-expressed with GFP-LC3. Scale bar, 10 µm. **d**, Per-cell quantification of Halo-WIPI2 membrane-tethering foci (top, red), GFP-LC3 foci (middle, green), and WIPI2^+^/LC3^+^ colocalized foci (bottom, yellow). **e**,**f**, Per-event intensity analysis for ATG13 (**e**) and GFP-LC3 (**f**) tracks across all conditions. Three intensity descriptors (peak, initial, and mean) are reported for the full track population (“all tracks”), and paired peak intensities are reported for ATG13^+^/LC3^+^ colocalized events (“colocalized”). Heatmap color encodes log_2_ fold change relative to the vehicle Fed condition; blue, decreased; red, increased. Boxplots in **b**,**d** show median (line), mean (open square), interquartile range (box), and 1SD (whiskers); individual cells are overlaid as gray points. Three biological replicates per condition (n = ≥80 cells per condition total). Statistical comparisons in **b**,**d** by ANOVA; *P < 0.05, **P < 0.01, ***P < 0.001, ****P < 0.0001. Statistical comparisons in **e**,**f** are versus the vehicle-Fed condition by Welch’s t-test on per-replicate medians with Holm–Bonferroni correction across the displayed comparisons; *P < 0.05, **P < 0.01, ***P < 0.001.

ATG13 initiation foci were largely preserved across the metformin dose range. Foci numbers and Baf-induced accumulation remained comparable to vehicle controls at all three doses tested (**Fig. 2b, top; Movies S1-4**). The autophagic initiation signal – recruitment of the ULK1-ATG13-FIP200 complex to nascent assembly sites – was therefore not interrupted by metformin, regardless of concentration. This finding is consistent with previous reports that metformin’s effect on mTORC1 is dose-dependent and occurs only at supratherapeutic concentrations ^21^ and indicates that the autophagic defect must lie downstream of initiation. WIPI2 foci, by contrast, decreased in a dose-dependent manner. The reduction was already significant at the lowest, clinically relevant 5 µM dose (p < 0.0001), reached its maximum at 200 µM, and persisted at 500 µM (**Fig. 2d, top; Movies S5-8**). The dissociation between intact ATG13 initiation and collapsed WIPI2 tethering localizes metformin’s molecular point of action to the membrane-tethering step that links the upstream initiation complex to nascent autophagosomal membranes. We previously identified this same step as the substrate of AMPK-driven autophagy suppression in glucose starvation, ^16^ where impaired ATG9-vesicle tethering by WIPI2 generated short-lived, abortive phagophores. The metformin phenotype, therefore, engages the same molecular bottleneck through a pharmacologically distinct upstream input.

This tethering defect translated directly into an LC3 flux deficit. GFP-LC3 puncta still accumulated under Baf in metformin-treated cells across all doses (**Fig. 2b, d, middle**), recapitulating the LC3-II accumulation interpreted as autophagy induction across the metformin literature. However, the LC3 foci that colocalized with ATG13 or WIPI2 – the population that reports on productive autophagosome assembly rather than residual lipidation events – were sharply reduced under metformin in both cell lines (**Fig. 2b, d, bottom**). Quantitative analysis of per-event foci intensities supported this finding: paired LC3 peak intensities at ATG13-positive structures were significantly attenuated across all metformin conditions (**Fig. 2e–f**, paired LC3 peak columns), with the effect strongest at 5 µM and remaining significant up to 500 µM. Initial and mean intensities of ATG13 tracks were similarly reduced (**Fig. 1e**, all-tracks columns), indicating that even when initiation foci form, they recruit less LC3 over their lifetime. Together, these per-event signatures diagnose autophagosomes that initiate but fail to complete membrane expansion, the molecular hallmark of abortive autophagy. ^4^ The dissociation between bulk LC3 accumulation, which appears induced under Baf, and productive colocalized flux, which is suppressed, directly explains the contradictory conclusions drawn from static LC3-II readouts across the metformin literature.

We next asked whether direct AMPK activation could substitute for, or rescue, metformin’s effect. Treatment with O-304 alone reduced productive LC3 colocalization in vehicle-treated cells, reproducing the maturation phenotype in wildtype AMPK cells (**Fig. 1b and d, bottom**). However, when O-304 was added on top of metformin, it produced no further reduction beyond metformin alone (rescue test, p > 0.05): the two perturbations converge at the maturation phase, consistent with a shared downstream effector. Critically, O-304 alone did not reproduce the dose-dependent dispersal of WIPI2 foci that defined metformin’s molecular signature: WIPI2 foci numbers in O-304-treated cells remained comparable to vehicle controls (**Fig. 1d, top**). AMPK activation can therefore account for the maturation phenotype but does not by itself drive WIPI2 displacement. Finally, we asked whether the reduction of LC3 foci was metformin-specific or shared by other complex I inhibitors. To test this, we used rotenone, which at low nanomolar concentrations selectively inhibits complex I. ^22^ Rotenone reproduced the metformin phenotype – a decrease in LC3 and WIPI2 puncta **(Fig. S1)** – indicating that the observed autophagy suppression is not metformin-specific but a general consequence of complex I inhibition. Taken together, these findings establish three points that frame the rest of the paper. First, complex I inhibition uncouples productive autophagosome assembly from bulk LC3 lipidation. Second, the decoupling is selective: initiation is preserved, tethering is lost. Third, the maturation phenotype is shared with direct AMPK activation. Still, the upstream WIPI2-tethering defect is not, suggesting that complex I inhibition engages a tethering input beyond what AMPK activation alone can deliver, which we next dissect kinetically and genetically.

### Complex I inhibition shortens autophagic foci dwell time and selectively displaces ATG13 and WIPI2 from membrane-tethered states

To define the kinetic basis of complex I inhibition effects on autophagy, we measured how long individual autophagic foci persisted before disappearing, i.e., the dwell time of ATG13-positive foci (a readout of initiation) and of LC3-positive foci (a readout of maturation).

Because foci still present at the end of imaging cannot be assigned a complete lifetime, we analyzed these distributions using survival statistics designed to handle such incomplete observations. Foci dwell-time distributions are summarized as Kaplan-Meier curves (**Fig. 3a**), and treatment effects are quantified by Cox proportional-hazards regression. A hazard ratio (HR) greater than 1 indicates faster focus dissolution – that is, shorter dwell times – under treatment relative to control. Inhibition of complex I with metformin progressively shortened LC3 dwell times across the entire tested dose range. KM curves showed that fed cells maintained a longlived foci population that was lost at all metformin concentrations (**Fig. 3a, top**). Cox regression confirmed similar effect sizes from 5 µM through 500 µM: HR = 1.37 (95 % CI 1.29–1.45, p < 10^−25^) at 5 µM, HR = 1.49 (1.28–1.74, p = 2 × 10^−7^) at 200 µM, and HR = 1.44 (1.32–1.58, p < 10^−15^) at 500 µM (**Fig. 3b**, right). The near-identical hazard ratios across a 100-fold concentration range indicate that the LC3 phenotype saturates at the low end of the metformin dose range. ATG13 dwell times tracked metformin similarly at low and intermediate doses (HR = 1.21 at 5 µM and 1.30 at 200 µM, both p < 0.001) but were not significantly affected at 500 µM (HR = 1.12, p = 0.16; **Fig. 3b**, left), consistent with a partial divergence between initiation and maturation phenotypes at high metformin concentrations.

**Fig. 3.**
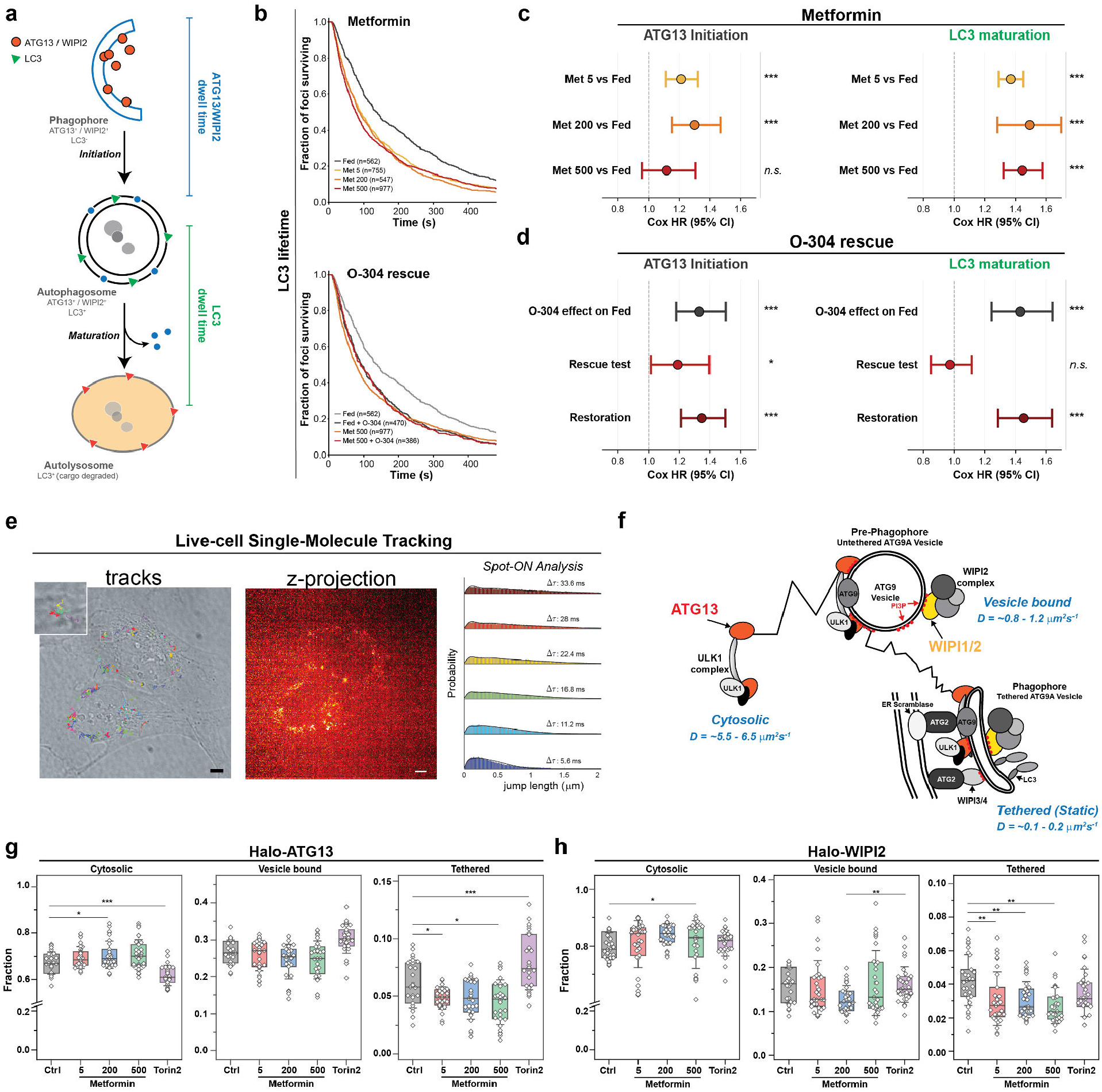
Metformin accelerates autophagic foci turnover and displaces WIPI2 from membrane-tethered states. **a**, Schematic of the autophagic intermediates and the dwell-time intervals measured in this study (left): the ATG13/WIPI2 dwell time on phagophores (initiation; LC3^−^), and the LC3 dwell time on maturing autophagosomes (LC3^+^). **b**, Kaplan-Meier survival curves of LC3 foci lifetimes in cells treated with vehicle (Fed) or metformin (5, 200, or 500 µM; top), and in cells treated with the direct AMPK activator O-304 (1 µM) alone or in combination with 500 µM metformin (bottom). Right-censored foci surviving to the end of the imaging window are included in the survival estimate (see Methods). Track counts are indicated. **c**, Cox proportional-hazards estimates for ATG13 initiation dwell time (left) and LC3 maturation dwell time (right) under each metformin dose versus the Fed condition. **d**, Cox proportional-hazards estimates for the O-304 rescue series: O-304 effect on Fed cells (top), rescue test (500 µM metformin + O-304 vs. 500 µM metformin alone; middle), and restoration test (500 µM metformin + O-304 vs. Fed; bottom). Hazard ratios > 1 indicate accelerated focus dissolution and shorter dwell times. Whiskers indicate 95% cluster-robust confidence intervals, with biological replicate as the clustering variable. **e**, Single-molecule tracking workflow: representative tracks overlaid on the brightfield image (left; inset 5× magnified), time-projected fluorescence image (middle; scale bar, 5 µm), and SpotOn jump-length distributions across multiple time lags (right) used to fit a three-state diffusion model. **f**, Schematic of the three molecular states resolved by SpotOn analysis: a fast cytosolic state, an intermediate vesicle-bound state corresponding to ATG9-vesicle association, and a slow membrane-tethered state at sites of autophagosome assembly. **g**,**h**, State occupancies for Halo-ATG13 (**g**) and Halo-WIPI2 (**h**) across conditions: vehicle (Ctrl), metformin (5, 200, 500 µM), and Torin2 (1µM), shown as cytosolic, vesicle-bound, and tethered fractions per cell. Boxplots in **g**,**h** show median (line), interquartile range (box), and 1SD (whiskers); individual cells overlaid as gray diamonds. Statistical comparisons in **g**,**h** are versus the Ctrl condition by ANOVA; *P < 0.05, **P < 0.01, ***P < 0.001. Each condition was sampled across three independent biological replicates with ≥ 25 cells per replicate per protein.

To test whether direct AMPK activation could substitute for, or rescue, the metformin effect, we performed the dwell analysis on the O-304-treated cells (**Fig. 3a**, bottom). O-304 alone shortened LC3 dwell times to a degree indistinguishable from metformin (HR = 1.43 vs fed, 1.25–1.64, p = 4 × 10^-7^), and the metformin + O-304 combination overlapped completely with metformin alone, returning a null rescue test (HR = 0.97, 0.85–1.12, p = 0.69; **Fig. 3c**, right). Despite the absence of additivity, the combination remained as different from the fed state as metformin alone (restoration test HR = 1.45, p = 2 × 10^-9^), confirming that O-304 does not relieve the metformin phenotype but rather phenocopies it, consistent with the two perturbations converging on a common downstream effector at the maturation phase. The ATG13 channel resolved a complementary asymmetry: O-304 alone shortened ATG13 dwell times (p < 0.001), and the rescue test was modestly significant (HR = 1.19, p < 0.05; **Fig. 3c**, left), indicating an additive rather than convergent effect at the initiation phase. The two pharmacological interventions therefore engage the same effector at LC3 maturation but separable inputs at ATG13 initiation.

To resolve the molecular state changes underlying the kinetic phenotype, we applied single-molecule tracking to endogenous Halo-ATG13 and Halo-WIPI2 (**Fig. 3d**). ^4^ This approach reports the diffusion of individual protein molecules, with mobility set by what each molecule is bound to: freely diffusing cytosolic molecules move fastest; in contrast, those engaged with vesicles or tethered to membranes move progressively more slowly. Trajectories partitioned into three diffusion states (Fig. 3e): a fast cytosolic state (D ≈ 5.5– 6.5 µm^2^ s^−1^), an intermediate vesicle-bound state (D ≈ 0.8– 1.2 µm^2^ s^−1^), and a slow tethered state (D ≈ 0.1–0.2 µm^2^ s^−1^) corresponding to membrane-engaged molecules at sites of autophagosome assembly. ^4^ Metformin produced a dosedependent loss of WIPI2 from the tethered state (p < 0.01 at 200 µM and p < 0.001 at 500 µM) with reciprocal increases in the cytosolic fraction (**Fig. 3g**). ATG13 was largely preserved across the metformin dose range, with only modest effects on the tethered fraction at 5 and 500 µM (**Fig. 3f**). The Torin2 control produced the inverse signature: a marked accumulation of ATG13 in the tethered state (p < 0.001) without a comparable redistribution of WIPI2 (**Fig. 3f, g**). This dissociation provides a molecular explanation for the kinetic phenotype: under metformin, autophagic structures initiate but fail to recruit the WIPI2-mediated tethering machinery required for ATG9-vesicle delivery and membrane expansion, generating short-lived, abortive foci. ^16^ Torin2, by contrast, accumulates ATG13 at autophagic structures consistent with canonical initiation, leaving the WIPI2-membrane interface intact.

### Restoring cytosolic NAD^+^, but not electron flux, prevents the metformin-induced loss of LC3 puncta

Complex I inhibition lowers both the NAD^+^/NADH ratio and the adenylate ratio, so metformin alone cannot distinguish which output drives the autophagy phenotype. Two reagents were compared: α-ketobutyrate (α-KB, 0.5mM), which is reduced by cytosolic lactate dehydrogenase and therefore regenerates NAD^+^ without supplying respiratory substrate, ^23^ and dimethyl succinate (DMS, 1mM), which feeds electrons to Complex II and restores Q-flux without consuming NADH (**Fig. 4a**). ^24,25^ Because electron input at Complex II reduces the ubiquinone pool almost completely, ^26^ succinate also depletes the oxidized ubiquinone that Complex I requires as an electron acceptor, so restoring Q-flux does not relieve, and can deepen, the redox lesion. Before interpreting the imaging experiments, we confirmed that each reagent corrected its intended arm, measuring both after 2 h of treatment (**Fig. 4b**). At 50 µM metformin, α-ketobutyrate returned NAD^+^/NADH to 7.80 and ATP/ADP to 2.48, recovering 31% and 10% of the respective deficits. Dimethyl succinate returned ATP/ADP to 2.69, recovering 29% of that deficit, and lowered NAD^+^/NADH further, to 5.83. Each reagent therefore corrected the arm it was selected for and left the other uncorrected or worse. These measurements are of whole-cell extracts and do not resolve the cytosolic and mitochondrial NAD pools. Because α-KB acts through cytosolic lactate dehydrogenase, it should correct the cytosolic pool while leaving the mitochondrial pool unchanged, and a whole-cell measurement averaging the two would report the partial correction observed here.

**Fig. 4.**
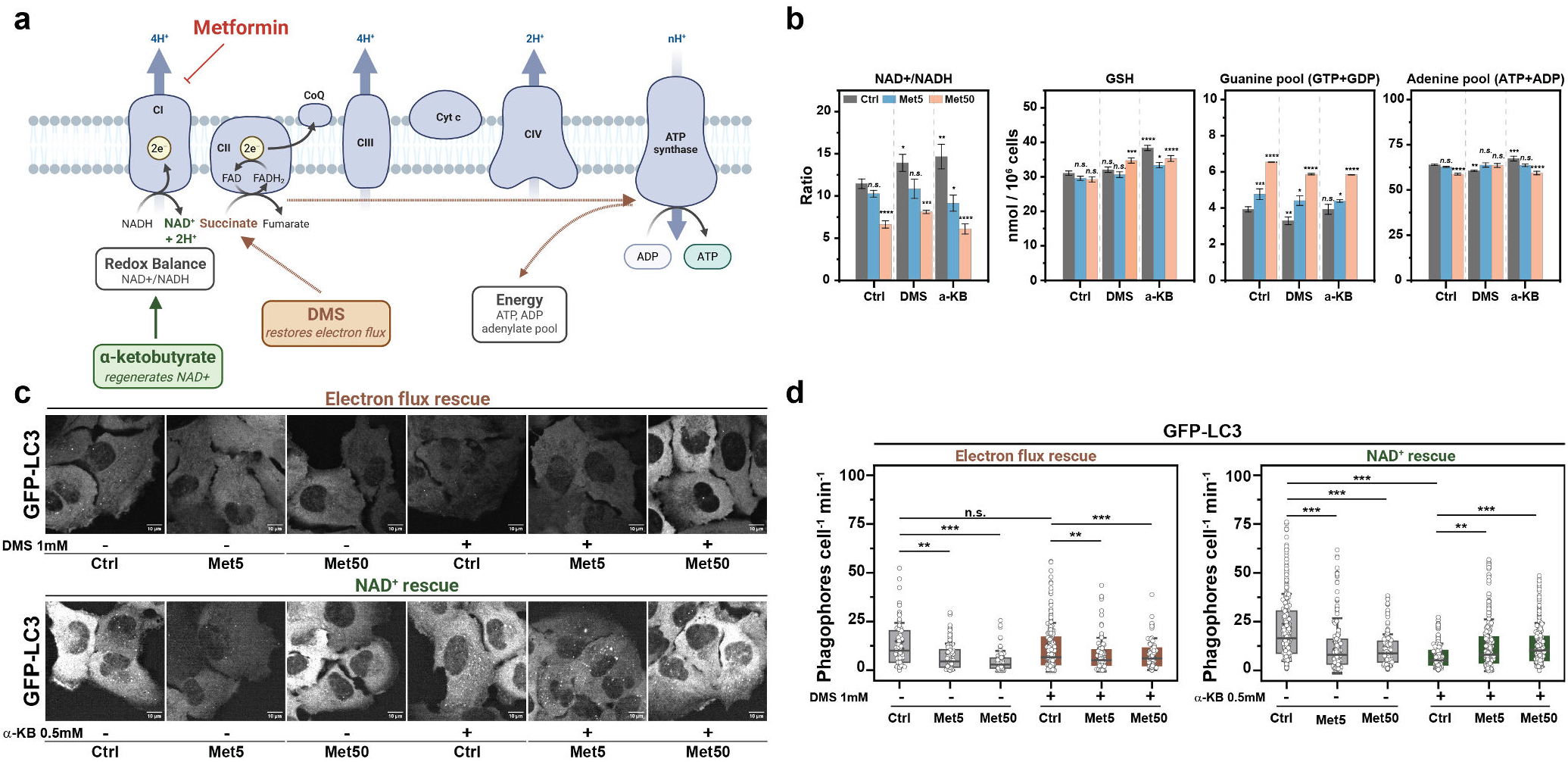
Restoring NAD^+^, but not electron flux, preserves phagophore formation under metformin. **a**, Schematic of the mitochondrial electron transport chain showing the two metabolic outputs of Complex I inhibition and the reagents used to separate them. Metformin inhibits Complex I, which simultaneously blocks NADH reoxidation (redox arm) and electron delivery to the Q pool (bioenergetic arm). α-ketobutyrate (0.5 mM) is reduced by cytosolic lactate dehydrogenase, regenerating NAD^+^ without supplying respiratory substrate. Dimethyl succinate (1 mM) donates electrons at Complex II, restoring Q-pool reduction and downstream proton pumping without consuming NADH. Created with Biorender.com. **b**, HPLC quantification of NAD^+^/NADH, glutathione, total guanine nucleotides (GTP+GDP) and total adenine nucleotides (ATP+ADP) in U2OS cells after 2 h of treatment, in the absence of a rescue reagent (Ctrl) or in the presence of dimethyl succinate (DMS) or α-ketobutyrate (α-KB). Bars are mean ± SD of three independent biological replicates performed on separate days; concentrations are nmol per 10^6^ cells and ratios are computed within each replicate before averaging. *P* values are Dunnett tests against untreated control (n.s., not significant; \**P* < 0.05; \*\**P* < 0.01; \*\*\**P* < 0.001; \*\*\*\**P* < 0.0001). **c**, Representative single frames of live U2OS cells expressing GFP-LC3 after 2 h of treatment with metformin (5 or 50 µM) alone or together with dimethyl succinate (top) or α-ketobutyrate (bottom). Scale bars, 10 µm. **d**, Phagophore formation rate for the electron-flux rescue (left, DMS 1 mM) and the NAD^+^ rescue (right, α-KB 0.5 mM). Boxes show the median and interquartile range, whiskers the 10th–90th percentiles. *P* values are from ANOVA (\*\**P* < 0.01; \*\*\**P* < 0.001).

To define whether the metformin-induced autophagy defect is driven by the redox arm of Complex I inhibition or by its bioenergetic arm, we quantified GFP-LC3 puncta in live U2OS cells, following a 2 h metformin treatment with or without a metabolic rescue reagent (**Fig. 4c, Movies S9-10**). Under these conditions, co-treatment with α-KB prevented the metformin-induced loss of LC3 puncta. Cells treated with metformin together with 0.5 mM α-KB contained more LC3 puncta than cells treated with metformin alone and/or treated with α-KB alone, indicating that the combination exceeded both single agents rather than reflecting their sum (**Fig. 4c,d**). The number of foci in the co-treated cells was not significantly different from the untreated control, indicating restoration rather than partial attenuation. Because puncta detection depends on fluorescence intensity, we asked whether the additional puncta in co-treated cells could be explained by brighter GFP-LC3 rather than by more structures. At 5 µM metformin, per-punctum GFP-LC3 intensity did not differ between metformin and metformin + α-KB (0.91 and 0.85 of control, P = 0.56, data not shown), while puncta number differed 2-fold (**Fig. 4d**), indicating that the additional puncta are additional structures. LC3 conjugation depends on ATP-dependent activation by ATG7, which may couple the amount of LC3 loaded per structure to the bioenergetic arm. Together, these data indicate that restoring NAD^+^ availability, but not electron entry at Complex II, is sufficient to preserve LC3 foci number under metformin, placing the foci-number response on the redox arm of Complex I inhibition. Per-structure LC3 content responded to the opposite reagent, indicating that the two arms act at different steps.

### PEN2 is dispensable for metformin’s effects on autophagy

The rescue experiments place the input at Complex I. A recent model proposes that low-dose metformin acts instead at the lysosome, where PEN2 relays v-ATPase inhibition to AMPK. ^9^ Because the two models predict different upstream events, we tested the lysosomal route directly. In that model, low-dose metformin acts on the lysosomal surface, where PEN2 is required to relay v-ATPase inhibition to AMPK activation; we therefore first asked whether metformin alters lysosomal abundance. First, live-cell LysoSensor imaging in GFP-LC3 U2OS cells revealed no change in the number of acidic lysosomal puncta across metformin concentrations (5, 200, 500 µM, 2 h), whereas bafilomycin A1 reduced LysoSensor-positive puncta in every condition (**Fig. S2a, b**), confirming that the assay detects perturbation of the acidic compartment and that metformin does not measurably expand or contract it at the doses and time points driving our autophagy phenotypes. Second, and more decisively, we generated two independent PEN2 knockout clones in Halo-WIPI2 cells, confirmed by loss of the PEN2 band on immunoblot (**Fig. S2c**). In both clones, EBSS and bafilomycin A1 induced AMPK activation (measured as Thr172 phosphorylation) and LC3 lipidation to levels indistinguishable from parental U2OS (**Fig. S2c**). The canonical AMPK response to nutrient stress and v-ATPase inhibition is therefore fully intact in the absence of PEN2, directly contrary to the requirement on which the Ma et al. model rests.

PEN2 loss disrupted baseline autophagy independently of metformin. Halo-WIPI2 phagophore initiation collapsed in PEN2 KO cells (*∼*2–3 events cell^-1^ min^−1^ vs. *∼*10 in parental; **Fig. S2d**), and GFP-LC3 redistributed from discrete puncta into large circular structures (**Fig. S2e**), a morphology consistent with the lysosomal dysfunction reported in γ-secretase-deficient cells. ^27^ Against this collapsed baseline, metformin produced no further change in WIPI2 dynamics (**Fig. S2d**), a loss of dynamic range rather than evidence of epistasis. Finally, we tested whether metformin induces bulk autophagy in MEFs over the extended time course reported by Ma et al. ^9^ Over 24 h at 5, 200, and 500 µM, p62 failed to decrease at any dose and instead accumulated at 200 and 500 µM, while LC3-II co-accumulated in parallel (**Fig. S2f**), a signature of impaired rather than induced autophagic flux. Together, these data establish that the lysosomal PEN2 axis is dispensable for canonical AMPK signaling in our system and is not the route through which metformin reshapes autophagy.

### AMPK is required for WIPI2 displacement but restrains the ATG13 response to metformin

To define the contribution of AMPK to metformin’s effects on autophagy initiation, we generated knockout (KO) lines targeting individual AMPK subunits in the Halo-ATG13 and Halo-WIPI2 reporter backgrounds (**Fig. 5**). Consistent with AMPK functioning as an obligate αβγ heterotrimer, ^28^ ablation of any single subunit destabilized the others: β1 KO reduced β2 and α1/α2 protein levels, β2 KO reduced β1 and α1/α2, and α1/α2 DKO reduced both β1 and β2 (**Fig. 5a,b**). Across the panel, the targeted subunit was undetectable. In contrast, the residual subunits were coordinately lowered. The same genotypes showed reduced phosphorylation of ULK1 at Ser555, a canonical AMPK substrate ^29^ (**Fig. 5b**). Ser555 phosphorylation was lost in β1 KO and severely reduced in α1/α2 DKO and β2 KO under both fed and EBSS-starved conditions; together with the loss of the AMPK subunits themselves, this establishes these lines as AMPK-signaling-null. In contrast, bafilomycin A1-trapped LC3-II accumulation in response to EBSS was present in every genotype and indistinguishable from wild-type in all three KO lines (**Fig. 5c;** KO versus wildtype, not significant), demonstrating that bulk autophagic flux proceeds independently of AMPK and that the core machinery for initiation, elongation, and lipidation remains competent. These lines therefore provided genetically defined backgrounds in which AMPK signaling is functionally null while autophagic flux remains operational.

**Fig. 5.**
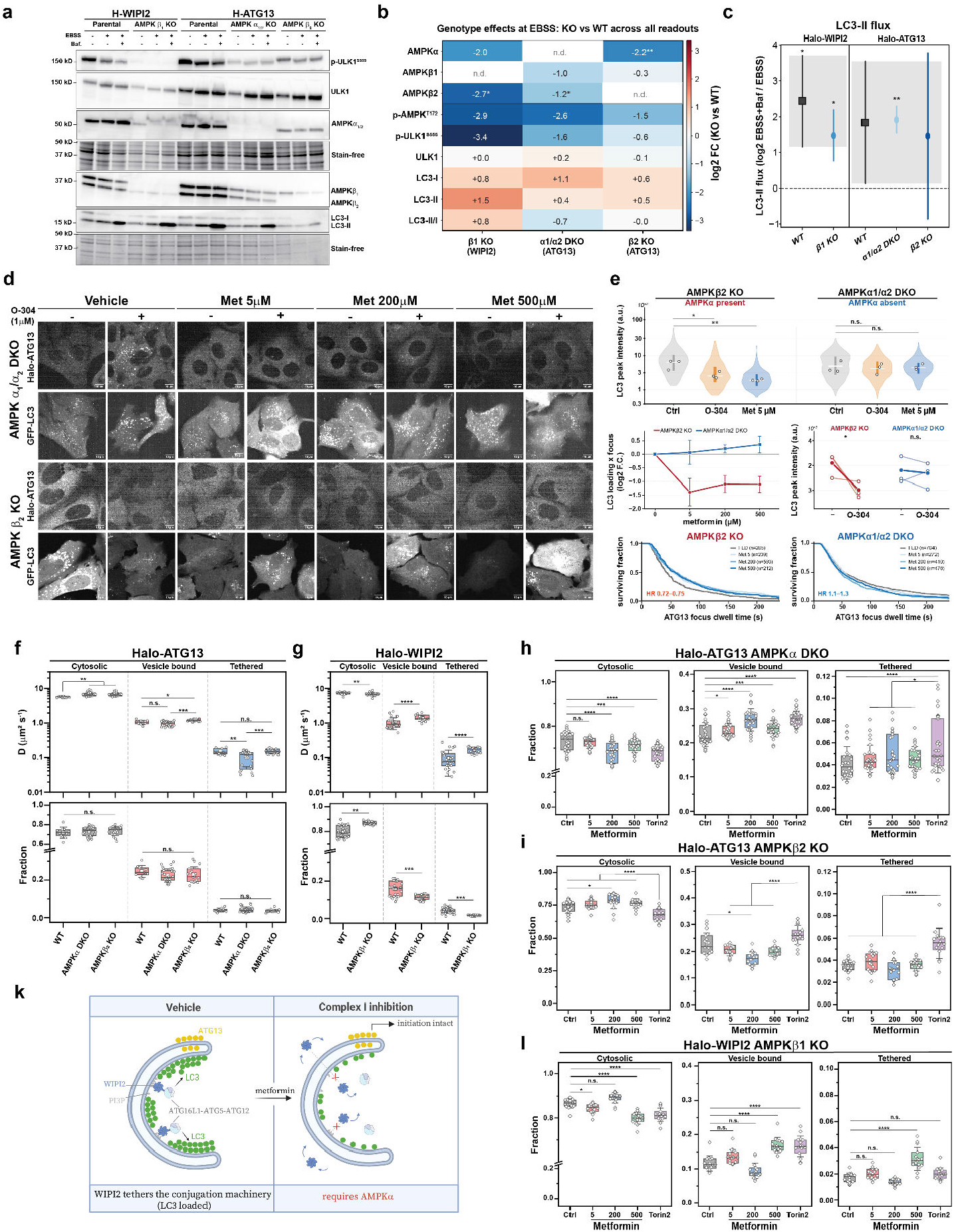
AMPK is dispensable for bulk autophagic flux but shapes the metformin response at initiation and LC3 conjugation in a subunit-specific manner. **a**, Immunoblots of Halo-WIPI2 (parental and AMPKβ1 KO) and Halo-ATG13 (parental, AMPKα1/α2 DKO and AMPKβ2 KO) in full medium or after 2 h EBSS, with or without 100 nM bafilomycin A1 (baf). Stain-free total protein serves as the loading control. **b**, Genotype effects under EBSS expressed as log_2_ fold change (KO versus the matched parental line) for each readout. Ablation of one AMPK subunit lowers the others, consistent with obligate heterotrimer assembly.: n.d., subunit not detected in the corresponding knockout. **c**, Bafilomycin-sensitive LC3-II flux, calculated as log_2_ (EBSS + bafilomycin / EBSS), for each genotype. Flux remains positive in every knockout, indicating that EBSS-induced bulk autophagy proceeds without AMPK. Grey bands denote the parental 95% confidence interval; points and error bars are mean ± 95% CI, n = 3. **d**,Representative live-cell confocal images of Halo-ATG13 and GFP-LC3 in AMPKα1/α2 DKO and AMPKβ2 KO cells after 2 h of vehicle or metformin (5, 200, 500 µM), with or without 1 µM O-304. Scale bars, 10 µm. **e**, Quantification of colocalized (ATG13^+^LC3^+^) foci. Top, LC3 peak intensity per focus; violins show the per-focus distribution, open circles are biological replicate medians. Middle left, metformin dose–response for LC3 loading; points are geometric means of replicate medians ± s.e.m. Middle right, paired per-replicate response to O-304. Bottom, Kaplan-Meier survival of ATG13 focus dwell time (1 s per frame; right-censored at the 239 s acquisition window); HR, hazard ratios from Cox proportional-hazards models with biological replicate as cluster. **f, g**, Baseline single-molecule behaviour of Halo-ATG13 (f) and Halo-WIPI2 (g) across genotypes, showing diffusion coefficients (top) and population fractions (bottom) for the cytosolic, vesicle-bound and tethered states. **h–j**, Metformin dose–response of state occupancy in AMPKα1/α2 DKO (h), AMPKβ2 KO (i) and AMPKβ1 KO (l) cells, with 1 µM Torin2 as a positive control. Redistribution between cytosolic, vesicle-bound and tethered pools is preserved in every AMPK-null background. Box plots show median, IQR, and whiskers = 1SD; each point is one cell from n = 3_ independent experiments. **K**, WIPI2 tethers the ATG16L1-ATG5-ATG12 conjugation machinery to PI3P on the phagophore, loading LC3; under Complex I inhibition WIPI2 is displaced and LC3 conjugation fails while ATG13 recruitment remains intact. For **e**, the statistical unit is the biological replicate; blocked ANOVA with replicate as block followed by Dunnett’s test against vehicle. n.s., *P* ≥ 0.05; \**P* < 0.05; \*\**P* < 0.01; \*\*\**P* < 0.001; \*\*\*\**P* < 0.0001. Created with Biorender.com.

To measure LC3 delivery to assembling autophagosomes in the same genetic backgrounds, we imaged Halo-ATG13 and GFP-LC3 in α1/α2 DKO and β2 KO cells after 2 h of vehicle, metformin (5, 200, 500 µM), or the direct AMPK activator O-304 (1 µM), and quantified peak LC3 intensity at ATG13-associated foci (**Fig. 5d,e, MoviesS11-12**). In β2 KO cells, metformin reduced LC3 loading per focus to 0.38-fold of vehicle at 5 µM (P = 0.004). The reduction was no deeper at 200 or 500 µM (0.47- and 0.46-fold; Met 5 versus Met 500, P = 0.13), so the effect was already maximal at the therapeutic concentration. The same reduction was present across the total LC3 focus population (0.46-to 0.50-fold across doses, P = 0.021), indicating that metformin lowers LC3 loading generally rather than only at ATG13-marked sites. In α1/α2 DKO cells, metformin produced no consistent change in LC3 loading. O-304 reproduced the reduction in β2 KO cells (0.48-fold, P = 0.029) and had no effect in α1/α2 DKO cells (0.93-fold, p = 1.000). Both metformin and O-304 therefore lower LC3 loading only in the genotype that retains AMPKα.

ATG13 residence at these foci diverged between the two genotypes. In β2 KO cells, metformin prolonged ATG13 dwell time across all three doses (HR 0.72–0.75 versus vehicle; **Fig. 5e**). In α1/α2 DKO cells, metformin shortened it (HR 1.1–1.3). ATG13 peak intensity was unchanged by metformin in both genotypes, so the difference in dwell time does not reflect a change in the amount of ATG13 recruited per site.

Single-molecule tracking under baseline conditions revealed that AMPK loss produces protein-specific biophysical phenotypes (**Fig. 5f,g**). In α1/α2 DKO cells, the tethered ATG13 population exhibited a marked reduction in its diffusion coefficient (D ≈ 0.08 vs 0.15 µm^2^ s^−1^ in parental; p < 0.01), indicating more confined membrane engagement without a measurable change in the fraction tethered (**Fig. 5f**). In β1 KO cells, WIPI2 was destabilized across all three diffusion states: diffusion coefficients were significantly elevated in each state, and the tethered and vesicle-bound fractions were reduced, with reciprocal accumulation in the cytosolic pool (**Fig. 5g**). Because ablation of any single subunit lowers the others (**Fig. 5a,b**), these phenotypes reflect loss of AMPK signaling rather than a function of the targeted subunit alone. Loss of AMPK signaling therefore alters the basal organization of the initiation machinery for both factors, confining residual ATG13 engagement while broadly destabilizing WIPI2 membrane association.

We then applied the same tracking framework to ATG13 across genotypes under vehicle, metformin (5, 200, 500 µM), and Torin2. The metformin response of ATG13 in parental cells was minimal, whereas Torin2 alone produced the canonical mTOR-inhibition signature of cytosolic decrease with reciprocal increase in vesicle-bound and tethered fractions (**Fig. 5d**). In α1/α2 DKO cells, metformin produced a dose-dependent redistribution of ATG13 from the cytosolic to the vesicle-bound state, accompanied by elevation of the bound-state diffusion coefficient (**Fig. 5h, Fig. S3a**), a response absent in parental cells. β2 KO cells did not reproduce this unmasking: ATG13 fractions were largely insensitive to metformin while the Torin2 signature was preserved (**Fig. 5i, Fig. S3b**). Both genotypes carry functionally null AMPK signaling (**Fig. 5a,b**), yet only α1/α2 DKO unmasks a metformin-driven ATG13 redistribution, indicating that the ATG13 response depends on the specific AMPK subunit ablated rather than on net AMPK signaling activity alone.

We next tracked WIPI2 across genotypes. In parental cells, metformin dose-dependently displaced WIPI2 from the tethered state, with reciprocal accumulation in the cytosolic pool (cf. **Fig. 3g**). In β1 KO cells, metformin produced no change in the tethered fraction at 5 or 200 µM, and only high-dose metformin (500 µM) altered it, in the opposite direction to parental cells (**Fig. 5l, Fig. S3c**). This line cannot test whether AMPK is required: baseline tethered WIPI2 is already reduced approximately twofold relative to parental (**Fig. 5g**), below the level metformin produces in parental cells, so the absence of a low- and mid-dose response is occluded by a floor rather than evidence of dependence. AMPK activation alone is not sufficient to reproduce the phenotype: the direct AMPK activator O-304 did not recapitulate metformin’s effect on WIPI2 foci in parental cells (**Fig. 2, Fig. 3**), nor on WIPI2 displacement here. Metformin’s displacement of WIPI2 therefore cannot be driven by AMPK activation alone, implicating a second, Complex I-coupled input at the maturation step. This contrasts with the initiation step, where AMPK restrains a metformin-driven ATG13 redistribution that becomes apparent only when AMPK is lost (**Fig. 5h**), establishing a non-canonical, subunit-specific role for AMPK across the two steps of autophagosome assembly that metformin engages (**Fig. 5k**).

### Metformin and glucose stress release the ULK1 complex from the cytoskeleton

To map how metformin engages the autophagy-initiation complex, we generated U2OS cells carrying a homozygous knock-in of a 3XFLAG-Halo tag at the endogenous *ULK1* locus (**Fig. S4, Fig. 6a**) and performed anti-FLAG immunoprecipitation-mass spectrometry across eight conditions: metformin at 5, 50, and 500 µM, the direct AMPK activator O-304, glucose withdrawal (-Glc), amino acid withdrawal (-AA), vehicle control, and a 3XFLAG-blank, sampled at 2 h in biological triplicate. After a stringent pulldown-ratio cutoff against the FLAG-blank, we quantified 183 ULK1 interactors across all conditions. The closest comparable prior work, an AP-MS/proximitylabelling interactome of all four ULK1-complex subunits under fed and starved conditions, ^30^ described the holo-complex as a signalosome but did not resolve a dose-response to a metabolic drug; our dataset captures, in a single coherent experiment, both the steady-state ULK1 interactome and the directional change driven by metformin and by orthogonal energetic stresses.

**Fig. 6.**
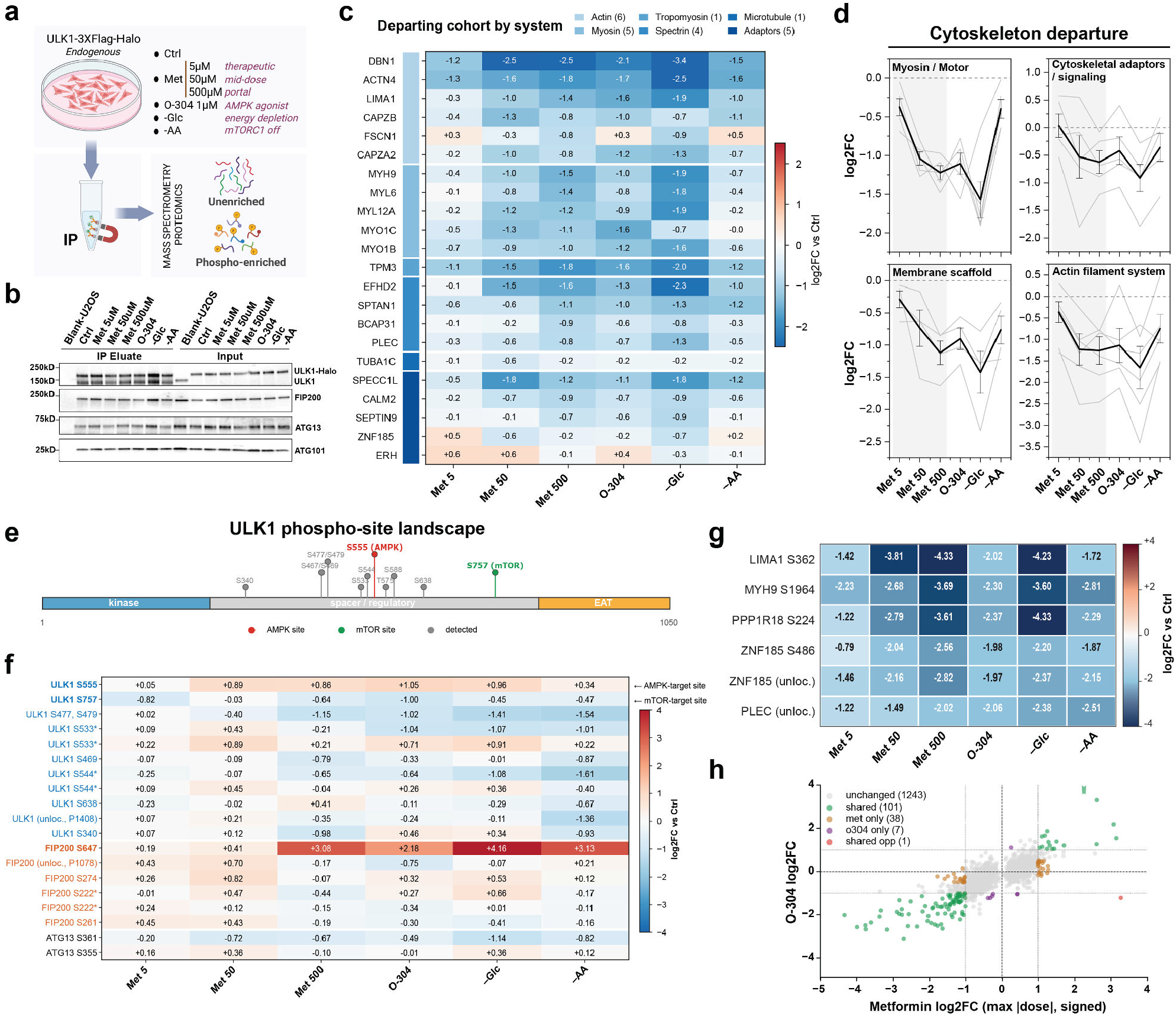
Metformin releases the ULK1 interactome from cytoskeletal scaffolds and restructures kinase inputs on the complex without altering its output on ATG13. **a**, Experimental design. U2OS cells carrying a homozygous knock-in of a 3×FLAG-Halo tag at the endogenous *ULK1* locus were treated for 2 h under seven conditions: vehicle (Ctrl), metformin at 5 µM (therapeutic), 50 µM (mid-dose) and 500 µM (portal), the direct AMPK activator O-304 (1 µM), glucose withdrawal (-Glc; energy depletion) and aminoacid withdrawal (-AA; mTORC1-inactivating). Anti-FLAG immunoprecipitates were analyzed in parallel by global proteomics (unenriched) and TiO_2_ /Fe-NTA phosphopeptide enrichment (phospho-enriched). Created with BioRender.com. **b**, Anti-FLAG immunoprecipitates probed for the core ULK1 holo-complex (ULK1, FIP200/RB1CC1, ATG13, ATG101); inputs at right, with the 3×FLAG-blank parental U2OS line as a specificity control. **c**, Departing cohort: 22 ULK1 interactors annotated to cytoskeletal systems (color-coded at top: actin, 6; myosin, 5; tropomyosin, 1; spectrin, 4; microtubule, 1; adaptors, 5), shown as per-condition log_2_ fold change versus Ctrl. **d**, The same cohort decomposed by subsystem (myosin/motor, cytoskeletal adaptors/signalling, spectrin/membrane scaffold, actin filament system). Grey lines, individual interactors; black line, subsystem mean ± s.e.m. **e**, Domain organization of human ULK1 (1,050 aa), N-terminal kinase domain, central spacer/regulatory region, C-terminal EAT domain, with all phosphosites detected in this dataset annotated above. The canonical AMPK site Ser555 (red) and mTORC1 site Ser757 (green) are highlighted; other detected sites in grey. **f**, Phosphosites within the ULK1-FIP200-ATG13 core across the six treatments versus Ctrl, grouped by subunit (ULK1, blue; FIP200, orange; ATG13, black). Bold labels mark Ser555, Ser757 and FIP200 Ser647. Sites labelled “unloc.” are phosphopeptides with ambiguous localization within the indicated region; asterisks mark sites quantified independently from two or more distinct phosphopeptides, with peptide-level identifiers in Supplementary Information. **g**, Phosphosites on cytoskeletal interactors that depart the pulldown **(c)** are coherently dephosphorylated across all active conditions, tracking the bulk pulldown ratios for the same proteins. **h**, Per-site comparison of metformin (*x* -axis; signed maximum |log_2_ fold change| across the three doses) against O-304 (*y* -axis; log_2_ fold change versus Ctrl) for all quantified phosphosites. Categories are defined by a |log_2_ FC| ≥ 1 threshold on each axis: unchanged in both (grey, n = 1,243), shared with concordant sign (green, n = 101), metformin-only (orange, n = 38), O-304-only (purple, n = 7), and shared with opposite sign (pink, n = 1). Heatmap colour encodes log_2_ fold change versus vehicle Ctrl: c, shared scale −2.5 to +2.5; f,g, shared scale −4 to +4; blue, decreased; red, increased. Per-condition values are medians across three biological replicates. Cohort assignment and thresholds for differential phosphorylation (|log_2_ ratio| ≥ 1, *P* ≤ 0.05, Proteome Discoverer background-based ANOVA) are given in Methods. The fully quantified interactome roster (n = 183 proteins) and complete phosphopeptide intensities, sequences and site-localization probabilities are provided as Supplementary Information.

The holo-complex itself and key regulators that engage it remained stably associated across conditions. ATG13, FIP200/RB1CC1, and ATG101 were detected in our pulldowns but were not significantly enriched over the U2OS parental IP-blank immunoprecipitation by filtering; their coimmunoprecipitation with ULK1 across all seven conditions, in treatment-invariant abundance, was confirmed by western blot (**Fig. 6b**). The dominant effect of metformin and metabolic stress was a coherent departure of cytoskeletal scaffolds from the ULK1 interactome (**Fig. 6c**). Eight actin-binding and intermediate-filament proteins – ACTN4, DBN1, EFHD2, LIMA1, MYH9, PLEC, PPP1R18, and ZNF185 – fell by log_2_ ratios of −1.0 to −1.4 across Met500, O-304, and starvation conditions, with the strongest departures driven by glucose withdrawal (ACTN4, DBN1, and EFHD2). The cohort behaved as a single signal: the dose-resolved metformin axis recapitulated the rank order of the starvation and AMPK-activator conditions (**Fig. 6d**). At Met5, the cohort showed modest, coherent disengagement (*∼*0.5 log_2_ units below control), with a substantially larger effect at Met50 and Met500 (*∼*1 to 1.5 log_2_ units). Although the actin cytoskeleton has been implicated in early autophagosome biogenesis, ^31^ that role was localized to the PI3K complex rather than to ULK1 itself, with no direct ULK1-actin contact reported; the proteins leaving the ULK1 interactome here are predominantly actin-binding scaffolds, consistent with the disengagement reflecting loss of indirect, scaffold-mediated baseline association rather than displacement of a direct ULK1-actin interaction. A few mitochondrial proteins showed small, nonspecific increases in ULK1 association under metformin, O-304, and starvation (**Fig. S4c**). Still, mitochondrial annotations were not significantly enriched under stringent filtering, and we do not interpret this as selective mitochondrial recruitment. ^32^

The AMPK α1 subunit PRKAA1 was recovered as a stable ULK1 interactor across all conditions, with a modest positive trend under active conditions (log_2_ ratio +0.16 to +0.40) and no depletion. The maintained – and if anything, slightly enhanced – AMPK-ULK1 association under metformin and energy stress is consistent with the stabilized inhibitory AMPKULK1 complex described by Park et al., ^14^ in which AMPK activation reinforces rather than releases its binding to ULK1. At the resolution of the endogenous interactome, metformin preserves this AMPK-bound core while selectively remodeling the peripheral, compartment-marker composition of the complex. The mTORC1 components MTOR and RPTOR were not recovered in the interactome roster, consistent with the canonical view that mTORC1 engages ULK1 transiently as a kinase-substrate rather than as a stable co-complex. ULK1 therefore retained its core complex partners, its cargo-recognition adaptors, and its AMPK association under every treatment; what changed across conditions was its peripheral composition. Together, these data describe an interactome-level signature on the ULK1 complex: at the 2 h initiation window, Complex I inhibition and metabolic stress drive a coordinated relocation of the ULK1 interactome – disengagement from actin-binding cytoskeletal scaffolds – while leaving the core ULK1-ATG13-FIP200-ATG101 assembly, its AMPK association, and its cargo-recognition machinery intact. We next assessed whether this compositional remodeling is matched by signaling changes within the complex itself by phosphopeptide enrichment of the same immunoprecipitates.

### Complex I inhibition restructures kinase inputs on the ULK1 complex

Phosphopeptide enrichment of the same anti-FLAG immunoprecipitates resolved how metformin signals through the initiation complex itself. Within the ULK1-ATG13-FIP200 core, regulated phosphosites are sorted unambiguously by upstream-kinase motif class (**Fig. 6e,f**). The single basophilic site in the core complex, the canonical AMPK substrate Ser555, trended upward across metformin doses, O-304, and glucose withdrawal, with the largest change under O-304. None of these comparisons reached significance, and we therefore treat the site as a consistent trend rather than a quantified response. The canonical mTORC1 repressive site Ser758 decreased broadly under metformin and metabolic stress (log_2_ ratio −0.5 to −1.0), while the dual mTORC1/AMPK site Ser638 was directionally invariant – the behavior expected when opposing inputs co-vary on a single residue. Two proline-directed ULK1 sites previously characterized as PBK/TOPK-deposited, ULK1-inhibitory marks (Ser469 and Ser533) likewise decreased, consistent with relief of a PBK-mediated restraint. ATG13 Ser355 – the direct ULK1 substrate and an established readout of ULK1 catalytic activity, abolished by ULK1 kinase inactivation and by pharmacological ULK1 inhibition ^30^ – was invariant across all eight conditions (|log_2_ ratio| ≤ 0.36). Reciprocal movement of canonical AMPK and mTORC1 sites in the absence of any change in ULK1’s substrate-directed output indicates that metformin re-poises the inputs to the complex without driving net activation of ULK1 toward its own complex partner.

Set against this broader downshift of proline-directed phosphorylation on ULK1, a single proline-directed site on the scaffold subunit moved sharply in the opposite direction. FIP200 Ser647 – within a Ser/Pro-rich patch spanning residues 642–653 – was the most strongly induced site in the dataset: baseline at Met5 and Met50 but elevated 8-to 18-fold at Met500, glucose withdrawal, and amino acid withdrawal **(Fig. 6f**). The direct AMPK activator O-304 produced no reproducible induction. The Ser647 motif (KASVSQTS647-PQSASSP) excludes both AMPK and ULK1 as direct kinases on consensus grounds (Pro at +1). An independent metformin phosphoproteomics study previously identified Ser647 and the adjacent Ser653 as metformin-regulated. ^33^ Substrate prediction (Kinase Library ^34^) and motif analysis place Ser647 in the proline-directed CMGC family, with stress-activated MAPKs (ERK7, JNK, p38) and atypical/transcriptional CDKs (CDK7, CDK8, CDK19, CDK17, CDK18) as the leading candidates. AMPK, mTOR, CK2, and ULK1 are excluded by motif (**Supplementary Table 3**). Notably, direct phosphorylation of the ULK1 complex by p38α MAPK at proline-directed sites has been demonstrated to inhibit ULK1 activity and disrupt the ULK1-ATG13 complex in inflammatory and metabolic contexts, ^35–37^ suggesting that a stress-activated MAPK input on FIP200 Ser647 may extend this regulatory axis to the scaffold subunit of the complex. The same prediction approach correctly identifies AMPK as the top candidate for ULK1 Ser556 (Supplementary Table 3), validating the method on a biochemically established target site.

Beyond the core complex, six phosphosites on actin-associated and cytoskeletal proteins (LIMA1, MYH9, PPP1R18, ZNF185, PLEC; **Fig. 6g**) showed strong, dosedependent loss across metformin and energy-stress conditions, paralleling the bulk pulldown departure of the broader cytoskeletal cohort (**Fig. 6d**) and arguing that these scaffolds leave the interactome rather than being retained in altered phosphorylation states. The breadth of this remodeling exceeded that of direct AMPK activation: at the phospho layer, metformin regulated 38 phosphosites not shared with O-304 against 7 O-304-exclusive sites, with 99% sign-concordance across the 102 shared events (Fig. 6h). Together, these data describe phospho-remodeling of the complex through at least three concurrent input axes – a basophilic AMPK input on Ser555, relief of a proline-directed restraint on Ser469/Ser533, and a proline-directed scaffold input on FIP200 Ser647 – concurrent with a falling mTORC1 input on Ser757 and an unchanged ULK1 substrate-directed output on ATG13 Ser355. At the 2 h initiation window, metformin’s inhibition of complex I rebalances multiple kinase inputs on the ULK1 complex while leaving its catalytic output toward ATG13 Ser355 unchanged. The kinase and functional consequences of FIP200 Ser647 phosphorylation, situated at the boundary between the ATG13-binding N-terminal domain and the membrane-binding coiled-coil, remain to be defined and are the subject of ongoing work.

### Complex I inhibition increases mitochondrial delivery to the lysosome

Complex I inhibition damages the mitochondria it acts on, suggesting that metformin should increase routing of mitochondria into the degradative pathway. We measured mitophagic delivery with a tandem reporter, cox8-GFP-mCherry, in which the cytochrome *c* oxidase subunit 8 presequence targets a GFP-mCherry fusion to mitochondria (**Fig. 7**). GFP fluorescence is quenched in the acidic lysosomal lumen whereas mCherry is not, so mitochondria delivered to the lysosome appear as mCherry-positive, GFP-negative puncta (red-only puncta). The reporter also revealed changes in mitochondrial morphology. Untreated cells contained an elongated, interconnected tubular network. Deferiprone fragmented this network into short, swollen mitochondria by 24 h, and CCCP produced fragmented mitochondria that collapsed into perinuclear clusters, with diffuse cytosolic reporter fluorescence. In metformin-treated cells the network remained largely tubular at both doses and timepoints (**Fig. 7a**), so the increase in lysosomal delivery occurred without extensive mitochondrial fragmentation. We scored red-only puncta per cell after 6 and 24 h of treatment. Metformin increased the number of red-only puncta per cell at both doses (**Fig. 7b**). At 6 h, the increase was modest (5 µM, 1.5; 50 µM, 1.3 red-only puncta per cell, versus 1.1 in untreated cells). By 24 h, both doses had approximately doubled the control value (5 µM, 1.9; 50 µM, 2.1). For every treatment, red-only puncta were more numerous at 24 h than at 6 h, so mitochondrial delivery to the lysosome accumulated over the treatment window rather than peaking by 6 h. The effect was present at the therapeutic 5 µM dose, not only at supratherapeutic concentrations. Metformin therefore induces mitophagy above the basal rate, and the magnitude of this induction remains within a low range rather than approaching the extensive mitochondrial clearance produced by uncouplers in Parkin-overexpressing cells. We included two mechanistically distinct inducers of mitophagy as positive controls. The iron chelator deferiprone induces mitophagy without requiring PINK1 stabilization or Parkin activation. ^38^

**Fig. 7.**
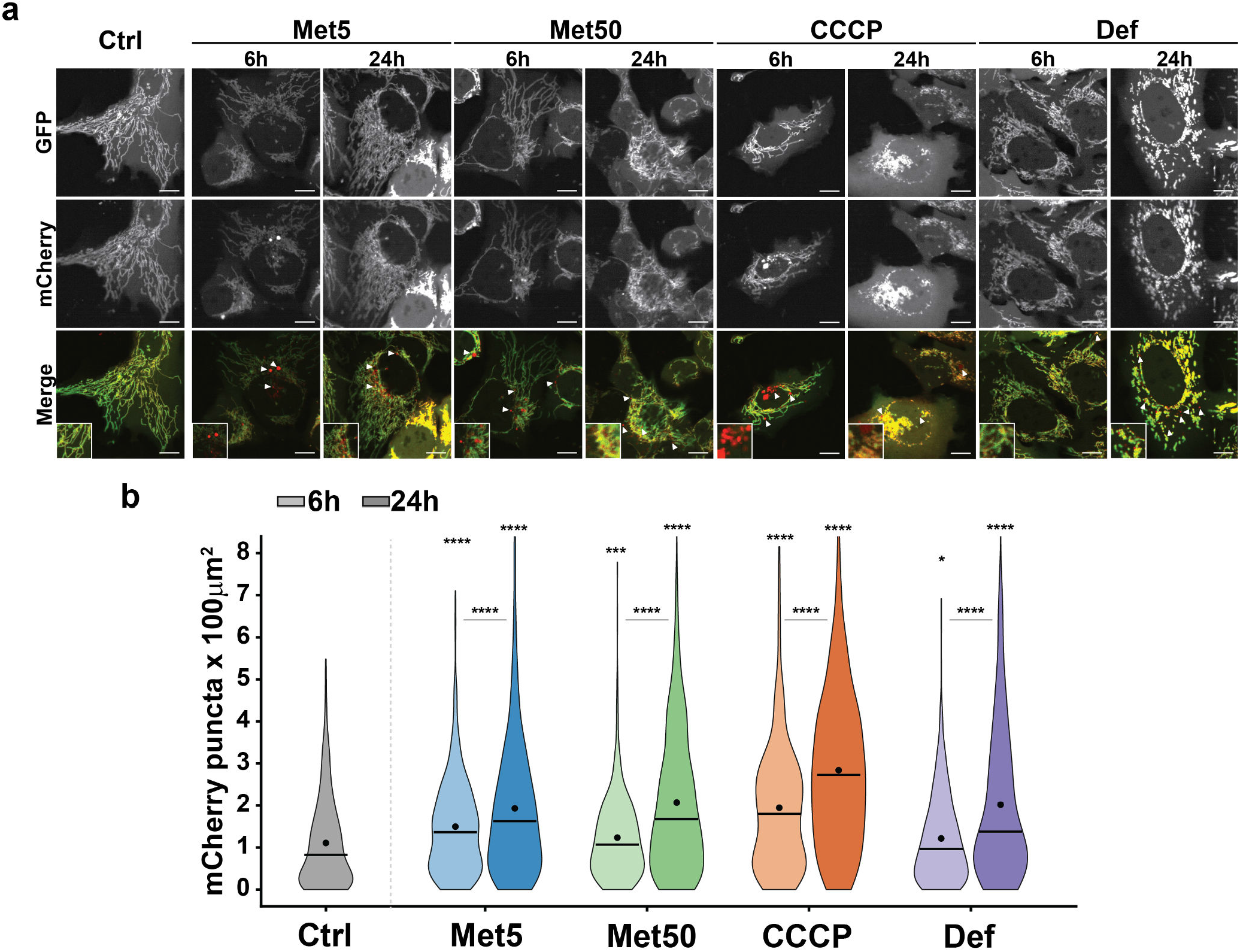
Metformin increases mitochondrial delivery to the lysosome. **a**, Representative confocal images of U2OS cells expressing the tandem mitophagy reporter cox8-GFP-mCherry, left untreated (Ctrl) or treated with metformin (5 or 50 µM), deferiprone (Def, 1 mM) or CCCP (1 µM) for 6 or 24 h. GFP and mCherry channels are shown individually and merged (GFP, green; mCherry, red). Red-only (mCherry^+^GFP^−^) puncta mark mitochondria delivered to acidic lysosomes, where GFP fluorescence is quenched. Insets show magnified regions. Scale bars, 10 µm. **b**, Quantification of red-only puncta per 100 µm^2^ of cell area. Violins show the distribution of per-cell values; horizontal bars indicate the median and dots the mean. Asterisks above each violin denote comparison with Ctrl; brackets denote comparison between 6 h and 24 h for each treatment. n > 150 cells per condition from three independent experiments. Mann-Whitney U test; *P < 0.05, ***P < 0.001, ****P < 0.0001.

Deferiprone raised the count at 24 h (*∼*2.0), and CCCP produced the largest increase at both time points (1.9 at 6 h, 2.8 at 24 h). Both controls confirm that the reporter registered lysosomal delivery of mitochondria in this cell background. U2OS cells express little Parkin. The response to deferiprone establishes that the assay detects Parkin-independent mitochondrial clearance, and basal mitophagy in mammalian tissues likewise proceeds independently of PINK1. ^39^ The metformin-driven increase in red-only puncta therefore does not require the canonical PINK1/Parkin pathway.

## Discussion

The major finding of this work is that pharmacological inhibition of complex I reprograms autophagy. At concentrations ranging from therapeutic to supratherapeutic levels, complex I inhibition generated autophagosomes that were short-lived, dim, and disconnected from WIPI2 (**Fig. 2, 3**). This signature reflects a block in maturation, not pathway induction. Direct AMPK activation with O-304 reproduced the same maturation phenotype, indicating that AMPK contributes to this step; however, O-304 did not reduce WIPI2 foci number, which remained comparable to vehicle controls. The result is consistent with a growing body of work in which AMPK engagement under metabolic stress restrains rather than promotes autophagy: glucose starvation, ^16^ sustained energy stress, ^14^ and prolonged amino-acid deprivation, ^15^ all converge on the maturation step of autophagosome assembly. Complex I inhibition reaches the same step from a mitochondrial input, and the effect is not specific to metformin: rotenone, at concentrations that selectively inhibit complex I, reproduced the decrease in LC3 and WIPI2 puncta.

Complex I inhibition perturbs cellular redox state before it perturbs the adenylate pool. In our metabolic time course, the NAD^+^/NADH ratio changed before any measurable change in the adenylate pool (**Fig. 1**). The same redox signal has recently been shown to engage AMPK directly through allosteric NAD^+^ binding at the γ1-CBS1 site, ^10^ providing a candidate route by which Complex I inhibition propagates to AMPK signaling in the absence of detectable changes in AMP/ATP and matching the AMPK engagement we observe at therapeutic metformin doses. Downstream of this metabolic input, the ULK1 holo-complex changes under metformin treatment: cytoskeletal scaffolds disengage, and a single proline-directed site on FIP200 (Ser647) is significantly induced, though only at supratherapeutic doses and under nutrient-deprived conditions. Ser647 induction is AMPK-independent: a direct pan-AMPK activator produced no reproducible induction, whereas Met500, glucose withdrawal, and amino-acid withdrawal each induced the site 8-to 18-fold. The Ser647 motif (KASVSQT-S647-PQSASSP) is consistent with this, excluding both AMPK and ULK1 as direct kinases on consensus grounds and identifying an unassigned proline-directed input to the complex, most plausibly within the CMGC group. The same site was previously flagged as metformin regulated. ^33^ The net effect of this reprogramming is neither suppression nor induction, but redirection (**Fig. 7–8**). Bulk cytosolic turnover is suppressed at the WIPI2-tethering step, while mitochondria are delivered to the lysosome, a substrate the bulk machinery no longer reaches (**Fig. 7**). Critically, this redirection occurs in U2OS cells, which lack detectable endogenous Parkin ^40^ and express the mitophagy receptors BNIP3 and NIX at low basal levels under standard culture conditions, making it unlikely that the canonical receptor-mediated mitophagy routes account for the clearance we observe and identifying a non-canonical mitochondrial clearance pathway engaged downstream of Complex I inhibition. The ULK1 complex therefore does not act as a simple switch between active and inactive autophagy. Its upstream metabolic state determines whether the machinery operates on cytosolic or mitochondrial cargo.

**Fig. 8.**
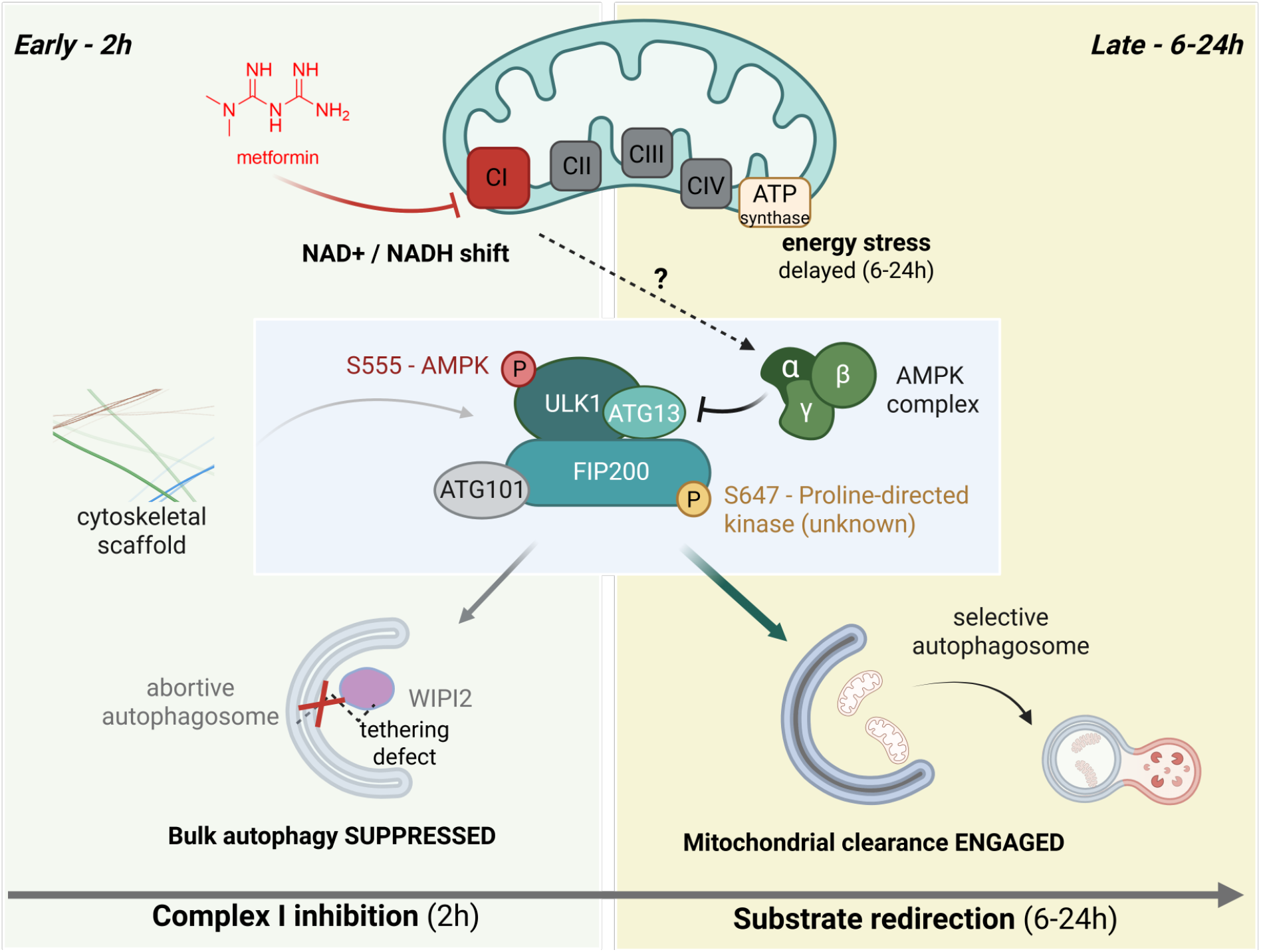
Complex I inhibition redirects autophagic substrate selection at the ULK1 complex. Working model integrating the metabolic, interactome, phosphoproteomic, and live-cell imaging data of this study. *Left (Early, 2 h):* Complex I inhibition raises the cellular NAD^+^/NADH ratio before the adenylate pool changes; adenylate depletion develop only at later time points (6–24 h) and are therefore not required for the tethering defect. Bulk cytosolic autophagy is suppressed at the WIPI2-tethering step: initiation proceeds normally while phagophores fail to mature, generating short-lived, abortive structures. *Center (ULK1 complex):* Complex I inhibition reprograms the composition and phosphorylation of the ULK1-ATG13-FIP200-ATG101 complex. Cytoskeletal scaffolds disengage in a dose-graded manner. AMPK remains stably associated and restrains the ATG13 response at initiation, while AMPK activation alone is not sufficient to reproduce the WIPI2 phenotype at maturation. An unidentified proline-directed kinase phosphorylates FIP200 Ser647 at supratherapeutic doses and under nutrient stress. *Right (Late, 6–24 h):* Bulk suppression persists while mitochondria are delivered to the lysosome, in Parkin-deficient cells, making canonical receptor-mediated mitophagy unlikely to account for the clearance observed. The ULK1 complex therefore does not act as a simple on/off switch: its upstream metabolic state determines whether the machinery operates on cytosolic or mitochondrial cargo. Created with Biorender.com.

Complex I inhibition reaches the autophagy machinery by a route distinct from the model currently dominant in the field. Genetic ablation of PEN2 in U2OS leaves the canonical AMPK response to nutrient stress and to v-ATPase inhibition intact, and metformin’s effects are not propagated through this lysosomal axis. AMPK is engaged downstream of Complex I, and its contribution differs between the two steps of autophagosome assembly (**Fig. 4**). At the maturation step, AMPK activation is not sufficient: O-304 did not reproduce the dose-dependent dispersal of WIPI2 foci, indicating that a Complex I-coupled input acts alongside AMPK. Whether AMPK is additionally required at this step cannot be determined from the AMPKβ1 KO line, because β1 loss by itself lowers the basal tethered WIPI2 fraction below the level metformin produces in parental cells (**Fig. 4c,g**), leaving no range in which a further displacement could be detected. That the signal converges on the WIPI2-tethering step is consistent with our previous report, using glucose starvation and pharmacological AMPK modulation, that AMPK gates phagophore-to-autophagosome maturation by controlling the tethering of WIPI2-positive phagophores to donor membranes. ^16^ The present genetic dissection identifies the same tethering step as the point at which a complex I-derived input is read out. AMPK suppresses the ATG13 response to complex I inhibition at initiation. In parental cells, metformin had little effect on ATG13 distribution. In AMPKα1/α2 DKO cells, the same doses produced a dose-dependent redistribution of ATG13 from the cytosolic to the vesicle-bound state (**Fig. 4e**). AMPKβ2 KO cells showed no such redistribution, although AMPK signaling is functionally null in both lines (**Fig. 4a**). The response therefore tracks which subunit is ablated, not the loss of AMPK activity itself. We are not aware of a previous report of AMPK suppressing an autophagy-initiation response in this way.

One question remains open: the identity of the proline-directed kinase acting at FIP200 Ser647. Candidate kinases, including stress MAPKs and transcriptional CDKs, are consistent with the motif, but assigning one requires direct genetic and chemical tests. Because the site responds only at supratherapeutic concentrations and under nutrient stress, identifying that kinase would also establish whether phosphorylation of the FIP200 scaffold contributes to substrate selection or reports a separate, higher-threshold response to energy stress. These results define a route by which a mitochondrial signal reaches the autophagy initiation machinery and changes what it degrades. Because metformin is taken chronically by a large patient population and has been associated with long-term metabolic and healthspan effects, ^41^ and because combination strategies pair it with bulk-flux inhibitors – metformin-gemcitabine in pancreatic cancer, for instance ^17^ – on the assumption that it acts as an autophagy inhibitor, the distinction between suppressing autophagy and redirecting it is likely to matter in both settings.

## Materials and Methods

### Chemicals

All chemicals and consumables were obtained from Thermo Fisher Scientific unless otherwise stated. Pharmacological agents: metformin hydrochloride (Cat. # AC135360010), the AMPK activator O-304 (Cayman Chemical, Cat. # 35828), the mitochondrial uncoupler CCCP (Cat. # AC228131000), the mTOR inhibitor Torin2 (Cat. # 50-115-0247), and the v-ATPase inhibitor bafilomycin A1 (Cat. # AAJ61835MCR). The HaloTag ligand JFX650 was provided as a gift from Luke D. Lavis (Janelia Research Campus). HPLC analytical standards (ATP, ADP, GTP, GDP, UTP, UDP, NAD^+^, NADH, reduced glutathione, sodium nitrite, and inosine), as well as dimethyl-succinate (DMS) and α-ketobutyrate, were obtained from Sigma-Aldrich. Ion-pairing reagents (tetrabutylammonium hydroxide, KH_2_PO_4_) and HPLC-grade methanol were from Thermo Fisher Scientific. Cell lysis and immunoprecipitation reagents included IP Lysis Buffer (Cat. # 87787) supplemented with 1× Halt Protease and Phosphatase Inhibitor Cocktail (Cat. # 78440), Pierce Anti-FLAG Magnetic Agarose (Cat. # A36797) for Halo-FLAG-ULK1 capture, and the Pierce High-Select TiO_2_ and Fe-NTA Phosphopeptide Enrichment Kits (Cat. # A32993) for SMOAC phosphopeptide enrichment. Mass spectrometry sample preparation reagents (TCEP, iodoacetamide, sequencing-grade trypsin, triethylammonium bicarbonate, calcium chloride, and formic acid) were obtained from Thermo Fisher Scientific. High-precision coverslips (170 ± 5 µm) for single-molecule imaging were from Thorlabs (Cat. # CG15CH).

### Antibodies

All primary antibodies were obtained from Cell Signaling Technology unless otherwise stated. The following antibodies were used for Western blotting at 1:1,000 dilution: anti-ULK1 (Cat. # 8054), anti-phospho-ULK1 Ser555 (Cat. # 5869), anti-phospho-ULK1 Ser757 (Cat. # 6888), anti-FIP200/RB1CC1 (Cat. # 12436), anti-ATG13 (Cat. # 13468), anti-ATG101 (Cat. # 13492), anti-AMPKα (pan, α1/α2; Cat. # 5831), anti-phospho-AMPKα Thr172 (Cat. # 2535), anti-AMPKβ1/β2 (Cat. # 4150), anti-LC3B (Cat. # 3868), anti-FLAG/DYKDDDDK tag (Cat. # 14793), anti-PEN2 (Cat. # 8502). HRP-conjugated secondary antibodies (goat anti-rabbit IgG, Cat. # 1706515; goat anti-mouse IgG, Cat. # 1706516) were from Bio-Rad and used at 1:2,000 dilution.

### Cell lines

All cell lines used in this study were derived from human bone osteosarcoma epithelial cells (U2OS, ATCC HTB-96). Mouse embryonic fibroblasts (MEF) were kindly donated by Joshua Andersen (University of Utah). We have extensively characterized the monoclonal cell lines edited to express HaloTag at the endogenous loci of the autophagy factors ATG13 and WIPI2. ^4^ ULK1-Halo was generated using the sgRNA (a gift from Jens Schmidt; Addgene plasmid # 207559; http://n2t.net/addgene:207559 ; RRID:Addgene_207559) and homology-recombinant plasmids (a gift from Jens Schmidt; Addgene plasmid # 207548; http://n2t.net/addgene:207548; RRID:Addgene_207548). ^4^ AMPK isoforms and PEN2 knockouts were generated by transfecting a plasmid encoding Cas9 and a single guide RNA (sgRNA, from VectorBuilder) targeting their respective loci into U2OS cells expressing Halo-ATG13 or Halo-WIPI2. We generated single-cell clones by FACS sorting using the GFP signal to select transfected cells. Knockouts were verified by western blotting. Cells were grown in RPMI cell culture media supplemented with 10% FBS, 100 U/ml penicillin, and 100 µg/ml streptomycin at 37°C with 5% CO_2_.

### Plasmid construction and genome editing

The GFPLC3B reporter was generated by cloning the coding sequences of LC3B into AAVS1-TRE3G-EGFP (AAVS1-TRE3G-EGFP was a gift from Su-Chun Zhang; plasmid #52343; Addgene; https://n2t.net/addgene:52343; RRID:Addgene_52343), including a TEV-protease cleavage site separating EGFP and LC3B. ^42^ GFP-tagged LC3 were stably expressed to parental HaloTagged cell lines (Halo-ATG13, Halo-WIPI2) and Halo-ATG13/Halo-WIPI2 knockouts (AMPKα_1/2_ KO, AMPKβ_1/2_ KO, PEN2 KO) by introducing the coding sequence at the AAVS1 safe-harbor locus (PPP1R12C) by cotransfection of the donor plasmid encoding tetracycline inducible GFP-LC3B and a plasmid encoding Cas9 and a sgRNA targeting the AAVS1 locus (AAVS1 T2 CRIPR in pX330 was a gift from Masato Kanemaki; plasmid #72833; Addgene; https://n2t.net/addgene:72833; RRID: Addgene_72833). ^43^ Polyclonal GFP-LC3B-expressing cell lines were obtained by selection with 1.0 µg/ml puromycin, followed by FACS by selecting cells lying within the 75^th^–95^th^ percentile of the GFP signal.

### Metabolic characterization

We simultaneously separated and quantified low-molecular-weight metabolites related to energy metabolism, mitochondrial activity, and oxidative/nitrosative stress biomarkers using ion-pairing HPLC. The analytical protocol used a binary step-gradient system optimized with tetrabutylammonium as the ion-pairing reagent (ThermoFisher Scientific). Two buffer solutions were used: Buffer A, consisting of 12 mM tetrabutylammonium hydroxide, 10 mM KH_2_PO_4_, and 10% methanol at pH 7.00, and Buffer B, containing 8 mM tetrabutylammonium hydroxide, 8 mM KH_2_PO_4_, and 40% methanol at pH 4.00. The gradient profile involved a controlled transition from 100% Buffer A to 0% Buffer A over a defined timeline, with specific intervals for concentration shifts. Chromatographic runs were conducted at a flow rate of 0.5 ml/min, with the column temperature maintained at 25°C. The HPLC system consisted of a Shimadzu apparatus coupled with a UV diode array detector (Shimadzu, SPD-M30A), equipped with a 5-cm light-path flow cell and set to acquire spectra between 190nm and 800nm. We acquired and analyzed data using Shimadzu software. We separated 12 low-molecular-weight metabolites on a Hypersil C-18 column (100 x 2.1 mm, 5µm particle size) with an inline pre-column. The analytes included high-energy phosphates (ATP, ADP, GTP, GDP, UTP, UDP,), oxidized and reduced nicotinic coenzymes (NAD^+^, NADH), reduced glutathione (GSH), nitrite (NO_2_), and Inosine. We identified and quantified compounds by retention time, absorption spectra, and peak areas relative to ultrapure standard mixtures. We quantified most metabolites at 260 nm, whereas GSH and nitrite were measured at 206 nm. In cell extracts, the total amount of proteins was determined according to the number of cells, and all concentration values of the aforementioned compounds were normalized to 1 million cells and expressed as nmol/million cells. The full metabolomic dataset is available in the supplementary information (Supplementary Data 1).

### Confocal live-cell imaging

We performed live-cell imaging experiments using an Olympus microscope (IX83). The microscope is equipped with an environmental chamber (cell-Vivo) to control humidity, temperature, and CO_2_ levels, a 60× total internal reflection fluorescence oil-immersion objective (Olympus UPlanApo, NA = 1.50), and appropriate excitation and emission filters. The microscope is equipped with two Hamamatsu Orca Fusion BT sCMOS cameras (Hamamatsu Photonics). We acquired images at 2×2 binning (1pxl = 0.217 µm). Cells were maintained at 37°C and 5% CO_2_ throughout imaging using a Tokai HIT stagetop incubation system. HaloTagged autophagy proteins labeled with JFX650 were excited using a Coherent OBIS 640 nm laser (80% power), while GFP-LC3 was imaged using a Coherent OBIS 488 nm laser (25% power). We acquired images at 1 frame per second with a 100 ms exposure time per channel. We imaged wild-type cells for 8 minutes (480 frames), whereas we imaged knockout cells for 4 minutes (240 frames). We quantified autophagy dynamics at the single-cell level using K-FOCUS, ^16^ a computational pipeline comprising three steps: single-cell segmentation, foci localization and tracking, and multimodal colocalization analysis. We segmented cells co-expressing endogenous HaloTagged ATG13 (CH1) and stably integrated GFP-LC3 (CH2) using CellPose 2.0, ^44^ a deep-learning–based tool trained on GFPLC3-expressing cells. We defined single-cell regions of interest (ROIs) on the GFP-LC3 channel (CH2) and applied them to the HaloTagged ATG13 channel (CH1) to ensure consistent cross-channel analysis. ROIs were imported into TrackIt for single-particle tracking using a nearest-neighbor algorithm with the following parameters: threshold factor of 1.2 for HaloTagged ATG13 (CH1) and 3.0 for GFP-LC3 (CH2); tracking radius of 4 pixels (*∼*0.870 µm); minimum track length of 5 frames; and gap tolerance of 5 frames. We assigned tracks to individual cells based on their first appearance within each ROI. Colocalization between ATG13 and GFP-LC3 foci was defined using the default K-FOCUS proximity criterion (≤3 pixels / 0.650 µm for ≥10 consecutive frames). We exported tracking data in batch format for downstream single-cell analysis.

### Lifetime analysis of autophagic foci

We imported tracklevel data, including per-frame coordinates, frame-by-frame intensity values, and a binary colocalization flag indicating spatiotemporal overlap between Halo-ATG13 or Halo-WIPI2 (CH1) and GFP-LC3 (CH2) foci, from the upstream K-FOCUS pipeline. We performed all subsequent analyses in Python (v.3.11); the analysis pipeline is provided as supplementary code.

#### Lifetime extraction and right-censoring

For each track, the focus lifetime was defined as the time elapsed between the first and last frames in which the focus was detected (frame interval = 1 s). Tracks whose final detection occurred at the last frame of the imaging window (480 s) were treated as right-censored, since their true lifetimes could not be observed within the experimental window. The fraction of right-censored tracks ranged from 1–7% across conditions. We preserved right-censoring in all survival analyses by encoding each track with a duration value and a binary event indicator (1 = observed dissolution, 0 = censored at the imaging cap), following standard survival-analysis practice. ^45,46^ Lifetime analyses reported in the main text were restricted to colocalized tracks (i.e., events in which ATG13 or WIPI2 and LC3 foci overlapped spatiotemporally as defined by the upstream colocalization pipeline) on the channel CH2 (LC3) population, capturing the dwell time of LC3 at autophagic foci.

#### Hierarchical data structure and unit of replication

Imagingderived foci data are inherently hierarchical: individual foci (tracks) are nested within cells, cells are nested within imaging sessions (biological replicates), and imaging sessions are nested within experimental conditions. Treating tracks as independent observations would constitute pseudoreplication, since tracks within the same cell share local membrane environment, drug exposure, and acquisition parameters, and cells within the same session share medium batch, focus, and imaging-day conditions. ^47^ To respect this nested structure, we performed primary statistical inferences at the biological-replicate level, using cell-level data as input to hierarchical models that explicitly account for between-replicate variability. For each (condition, biological replicate) combination, we imaged multiple cells (range: 23–78 cells per combination), yielding 80–160 cells per condition across three biological replicates. From each cell, a per-cell median lifetime was computed across all colocalized tracks belonging to that cell. Per-cell summary metrics also included the long-lived fraction (proportion of tracks with lifetime ≥ 60 s), and percell mean focus intensity, the latter providing an orthogonal readout of focus maturation state.

#### Statistical analysis

We applied two complementary statistical tiers to each pairwise comparison to triangulate the significance of the effect under different assumptions about the unit of replication.

Tier 1 (primary inference for figure annotation): a two-stage test in which cell-level metrics were first averaged within each biological replicate to yield one summary value per (condition, replicate) pair, followed by a paired Student’s *t*-test (when replicates overlapped between conditions) or Welch’s *t*-test (when they did not). This test treats the biological replicate as the unit of independence and is the most conservative of the three tiers.

Tier 2 (effect-size estimation for survival data): a Cox proportional-hazards regression on the track-level lifetime data with the biological replicate as a clustering variable, fit using the lifelines Python library. ^48^ Cluster-robust (sandwich) standard errors were used to correct for within-replicate correlation. ^49^ The Cox model returns hazard ratios (HRs) with 95% confidence intervals quantifying the relative rate of focus dissolution between conditions; an HR > 1 indicates faster turnover (shorter lifetimes).

Kaplan-Meier survival curves were used for descriptive visualization only and were not relied upon for inferential statistics. Pooled track-level log-rank tests were not used because they ignore the nested structure of the data and yield artificially small *p*-values. Tests were considered significant at *p* < 0.05 (two-sided). For any quantitative effect-size statement in the Results text, we used hazard ratios with cluster-robust 95% CIs from Tier 2. We performed all analyses in Python 3.11 with pandas, ^50^ NumPy, ^51^ SciPy, ^52^ statsmodels, ^53^ lifelines, ^48^ and matplotlib. ^54^ The complete analysis pipeline (.mat parser, cell-level aggregation, three-tier statistical framework, and figure-generation code) is available as supplementary code and will be deposited on GitHub upon publication.

### Single-molecule live-cell imaging

We characterized the diffusion behaviors of Halo-ATG13 and Halo-WIPI2 by single-particle tracking in living cells in response to vehicle (Ctrl), three doses of metformin (5, 200, and 500 µM), and Torin2. Imaging was performed on an Olympus microscope equipped with a 640 nm laser line (*∼*25% output) operating in HILO geometry and a 60×total internal reflection fluorescence oil-immersion objective (Olympus UPlanApo, NA = 1.50); emission was collected with an BT Fusion scMOS camera (Hamamatsu). For each experiment, 2–3 × 10^5^ cells were plated on high-precision coverslips (170 ± 5 µm, Thorlabs) and imaged 24 h post-seeding. We prepared coverslips as follows before mounting in 35-mm imaging dishes: 1 h sonication in 1 M KOH, rinsing in ddH_2_O, 1 h sonication in ethanol, and drying under an N_2_ stream. Sparse labeling was achieved using the JFX650 HaloTag ligand at 500 pM for 30 s, followed by media exchanges (3× wash). These conditions yielded 25–30 detectable particles per frame across the entire acquisition. Movies were recorded at 166 frames s^-1^ (5.6 ms exposure) over 2,000 frames within a 400 × 400-pixel ROI (1pxl = 0.217 µm). Drug-treated cells were imaged within 2 h of treatment onset. We performed particle localization and trajectory reconstruction in TrackIT using the nearest-neighbor algorithm. ^55^ Per-cell diffusion coefficients and population fractions were extracted from the resulting trajectories using the MATLAB implementation of SpotOn ^56^ (gift of the same authors) under a three-state kinetic model. SpotOn was configured as follows: TimeGap 5.6 ms, dZ 0.700 µm, GapsAllowed 2, Time-Points 8, JumpsToConsider 4, BinWidth 0.01 µm, PDF fitting; the diffusion bounds for the three states were D_Bound *∈* [10^-4^, 0.15], D_Free2 *∈* [0.15, 2], and D_Free1 *∈* [2, 15] µm^2^ s^-1^. We sampled each condition across three independent biological replicates with ≥20 cells per replicate per protein.

### Mitophagy quantification

The Cox8-GFP-mCherry reporter was generated by cloning the coding sequences of the cytochrome oxidase subunit 8 (Cox8) (a gift from Heather Wilkins, University of Kansas Medical Center) into AAVS1-TRE3G-EGFP (AAVS1-TRE3G-EGFP was a gift from Su-Chun Zhang; plasmid #52343; Addgene; https://n2t.net/addgene:52343; RRID:Addgene_52343) including a TEV-protease cleavage site separating EGFP and cox8. ^42^ GFP-mCherry-tagged Cox8 were stably expressed in U2OS by introducing the coding sequence at the AAVS1 safe-harbor locus (PPP1R12C) by cotransfection of the donor plasmid encoding tetracycline inducible Cox8-GFP-mCherry and a plasmid encoding Cas9 and a sgRNA targeting the AAVS1 locus (AAVS1 T2 CRIPR in pX330 was a gift from Masato Kanemaki; plasmid #72833; Addgene; https://n2t.net/addgene:72833; RRID: Addgene_72833). ^43^ Polyclonal cox8-GFP-mCherry-expressing cell lines were obtained by 2-week selection with 1.0 µg/ml puromycin.

Because mCherry is resistant to lysosomal acidification, whereas GFP fluorescence is quenched at low pH, delivery of mitochondria to the acidic lysosomal lumen yields mCherry-positive, GFP-low puncta (mitolysosomes), providing a ratiometric readout of mitophagic flux. Cells were treated in complete growth medium with metformin (5 or 50 µM), the mitochondrial uncoupler CCCP (1 µM, positive control), the iron-chelator deferiprone (Def, 1mM), or vehicle (untreated control) for a total of 6 h or 24 h. At the 6 h and 24 h endpoints, live cells were imaged on an Olympus IX83 inverted microscope equipped with a Yokogawa CSUW1 spinning-disk confocal unit, maintained at 37 °C and 5% CO_2_ using a Tokai HIT stage-top incubation system. We excited GFP with a Coherent OBIS 488 nm laser (30% laser power) and mCherry with a Coherent OBIS 561 nm laser (25% laser power), using identical exposure times (100 ms). Two-channel z-stacks (2304 × 2304 pxl, 0.1083 µm/pxl, 21 planes, 0.25 µm; mCherry then GFP) were maximum-intensity projected. Because we recorded the two channels on separate scMOS BT-Fusion cameras, we measured channel registration using bead calibration (TetraSpeck Micropheres 0.1µm, ThermoFisher Scientific). We segmented cells with Cellpose3 on the mCherry projection and manually curated the masks; we excluded cells smaller than 50 µm^2^, or those in which Otsu thresholding of the mCherry channel assigned less than 5% of the cell area to the mitochondrial network (1,474 cells from 77 images). Puncta were detected in two stages. A white *tophat* filter (disc radius 10 px, 1.08 µm) was applied to the mCherry projection, suppressing the tubular network, and candidates were thresholded at the median plus 8 robust standard deviations (1.4826 × MAD) of the in-cell *tophat* response. Candidates were retained at 8–800 px (0.09–9.4 µm^2^), eccentricity ≤ 0.90 and solidity ≥ 0.85. Each candidate was then normalized to the mitochondrial network of its own cell, defined by Otsu thresholding of mCherry within that cell: an mCherry+ punctum has a mean mCherry intensity ≥ 0.8 × the network median, and a mitolysosome is an mCherry+ punctum whose mean GFP is < 0.8 × the network GFP median, reflecting quenching of GFP in the acidified compartment. We set gate values on a subset of images spanning all conditions before analyzing the full dataset. We report both readouts as puncta per 100 µm^2^ of segmented cell area.

### ULK1 immunoprecipitation and proteomics

We used the ULK1-Halo cell line to perform immunoprecipitation proteomics and phosphorylation enrichment to define changes in the ULK1 complex upon metformin addition. We prepared complete RPMI media in the laboratory according to the standard protocol (Thermo Fisher Scientific). Glucose-free media (no glucose added) and amino acid-free media (no MEM amino acids and no L-glutamine added) were also prepared similarly. For all media prepared, we adjusted the pH to 7.3, filtered the media, and supplemented them with 10% fetal bovine serum (FBS) and 100 units/ml penicillin/streptomycin. To study the effects of metformin, we used various concentrations (5 µM, 50 µM, and 500 µM) as treatments. We also used the AMPK activator O-304 (1 µM). Control cells received only complete, prepared RPMI media, while test cells received complete media with either metformin or O-304; no-glucose cells received glucose-free media, and no-amino-acid cells received amino acid-free media for 2 hours before Immunoprecipitation (IP).

U2OS cells stably expressing ULK1-HaloTag were seeded in a 10 cm dish at 2 × 10^6^ cells on Day 0 and allowed to grow in RPMI cell culture medium supplemented with 10% fetal bovine serum (FBS) and 100 units/ml penicillin/streptomycin at 37°C with 5% CO_2_. On Day 1, the media was replaced with prepared complete RPMI media. On Day 2, the media were removed, and cells were washed twice with PBS before treatment. The cells were then incubated in the new media with or without treatment for 2 hours, as mentioned above. The media were then removed, and cells were washed twice with PBS before immunoprecipitation. PBS with 0.5 M EDTA was added to pre-washed cells and incubated at 37 °C for 10 minutes. The detached cells were then collected in a 15 mL conical tube and centrifuged at 300×g. The pellet was resuspended in 1 mL IP Lysis Buffer (Thermo Fisher Scientific), supplemented with 1x protease and phosphatase inhibitors (Thermo Fisher Scientific). The cell suspension was then passed through a 25G syringe to facilitate cell lysis. The suspension was then incubated for 10 minutes on ice with intermittent mixing. The lysed cells were then centrifuged at 13,000×g for 10 minutes. The Anti-Flag antibody beads (Thermo Fisher Scientific) were prepared for immunoprecipitation by washing 3× with PBS. The supernatant from lysed cells was added to the prepared beads, and the mixture was incubated at 4^0^C overnight for 16 hours. The next day, the beads were collected with a magnet and washed 2× with PBS, followed by a wash with DI H_2_O. The protein was then eluted from the beads and analyzed by western blotting and mass spectrometry.

### Mass spectrometry proteomics and phosphoproteomics

PBS was removed from FLAG-IP samples using a magnetic rack, and beads were resuspended in 100 µL of 50 mM TEAB, 2 mM CaCl_2_ prior to reduction by adding 0.5 M TCEP to a final concentration of 5 mM, followed by incubation at 55 °C for 30 minutes. We alkylated reduced samples by adding 375 mM iodoacetamide to a final concentration of 10 mM, then incubated them in the dark at room temperature for 30 minutes. We digested proteins on beads by adding Trypsin (500 ng) and incubating overnight at 37 °C with shaking at 1000 RPM (Thermomixer, Eppendorf). We quenched the digestion by adding 10% formic acid to a final concentration of 1%. We enriched samples for phosphorylated peptides using the SMOAC method. ^57^ Briefly, digested samples were first enriched by the High Select™ Phosphopeptide Enrichment kit (ThermoFisher Scientific) following the manufacturer’s protocol. The flow-through was applied to the High Select™ Fe-NTA Phosphopeptide Enrichment Kit (ThermoFisher Scientific) following the manufacturer’s protocol. The flow-through after the second enrichment became the global, unenriched sample, and the elutes from both kits were pooled to generate the phosphopeptide-enriched fraction. Samples were centrifuged at 10,000 x g for 10 minutes to remove particulates, and the supernatant was transferred to a fresh tube and stored at −20 °C until mass spectrometry analysis. Peptide concentrations were measured using a NanoDrop spectrophotometer (Thermo Scientific) at 205 nm. Samples were injected using the Vanquish Neo (Thermo) nano-UPLC onto a C18 trap column (0.3 mm x 5 mm, 5 µm C18) using pressure loading. Peptides were eluted onto the separation column (Aurora Elite XT, 15 cm x 75µm, 1.7 µm C18 particle size, IonOptics) heated to 40 °C prior to ionization at the mass spectrometer. Briefly, peptides were loaded and washed for 5 minutes at a flow rate of 0.350 µL/min at 2% B (mobile phase A: 0.1% formic acid in water, mobile phase B: 80% ACN, 0.1% formic acid in water). Peptides were eluted over 100 minutes from 2–25% mobile phase B before ramping to 40% B in 20 min. The column was washed for 15 min at 100% B, then re-equilibrated at 2% B for the next injection. The nano-LC was directly interfaced with the Orbitrap Ascend Tribrid mass spectrometer (Thermo) equipped with a high field asymmetric ion mobility spectrometry (FAIMS) source. The data were collected by data dependent acquisition with the intact peptide detected in the Orbitrap at 120,000 resolving power from 375–1500 m/z. Peptides with charge +2–7 were selected for fragmentation by higher energy collision dissociation (HCD) at 28% NCE and were detected in the ion trap at rapid scan rate (global) or in the Orbitrap at 30,000 resolving power (enriched). Dynamic exclusion was set to 60s after one instance. The mass list was shared between the FAIMS compensation voltages. FAIMS voltages were set at −45 (1.4 s), −60 (1 s), −75 (0.6 s) CV for a total duty cycle time of 3s. We set source ionization to 1700 V and the ion transfer tube temperature to 305 °C. Raw files were searched against the human protein database downloaded from UniProt on 03-21-2025, and a common contaminants database with variable phosphorylation allowed on S, T, and Y residues (enriched) using SEQUEST in Proteome Discoverer 3.0. ^58^ We exported abundances, abundance ratios, and p-values to Microsoft Excel for further analysis. The mass spectrometry data have been deposited to MassIVE (https://massive.ucsd.edu/). The accession numbers for the data reported in this paper are MassIVE MSV000102034 (global) and MSV000102035 (phosphopeptide-enriched). We also submitted them to ProteomeXchange (PXD079242 (global); PXD079244 (phosphopeptide-enriched)).

### Computational analysis of proteomics data

We performed downstream analysis of both arms of the anti-Flag-ULK1 IP experiment in RStudio (v.2026.01.2+418) using two custom R Markdown pipelines, deposited as supplementary files. Both pipelines take Proteome Discoverer outputs as input. Both the bulk and phospho arms apply Proteome Discoverer’s contaminant flag at the cleanup step, and we further restricted proteins to those with ≥ 2 unique peptides. An in-pipeline CRAPome-style contaminant list (n = 67, covering cytokeratins, tubulins, cytoplasmic actins, myosins, HSPs, glycolytic enzymes, hnRNPs, CCT chaperonins, S100s and immunoglobulin contaminants; based on the CRAPome empirical contaminant repository ^59^) was applied. We generated inter-sample Pearson correlations on log2-transformed normalized abundances with corrplot. ^60^ A protein was flagged as a ULK1 interactor if it was significantly enriched over the untagged-cell Blank control (log_2_FC ≥ 1, Welch’s t-test p ≤ 0.05) in at least one of the seven Flag-IP conditions versus Blank. Per-phosphopeptide IP-vs-Blank enrichment was retained as an annotation column but not used as an inclusion gate, because PD’s normalized ratios systematically underdetect abundant species against a low-abundance bait. The full proteomic and phosphoproteomic dataset is available as supplemental information (Supplementary Data 2).

### Statistics

Statistical analyses not described in the assay-specific sections above were performed as follows. We compared normally distributed continuous variables across multiple conditions (metabolomics, foci count in fixed-condition comparisons) using one-way analysis of variance (ANOVA), with Dunnett’s post-hoc test for comparisons against a vehicle control or Tukey’s HSD for pairwise comparisons across all groups. Two-group comparisons were performed by Student’s *t*-test (paired or unpaired as appropriate) or Welch’s *t*-test where variances were unequal. Where appropriate, multiple-comparison correction was applied using the Holm-Bonferroni method, as noted in the corresponding figure legends. All statistical tests were two-sided. Throughout, we considered *p < 0*.*05* significant, with significance levels indicated as \**p* < 0.05, \*\**p* < 0.01, \*\*\**p* < 0.001, and \*\*\*\**p* < 0.0001 in figure annotations. Data processing, statistical analyses, and figure generation were performed in OriginPro (version 2024b; OriginLab, Northampton MA) for metabolomics, foci-count comparisons, and routine plotting; and in R (version 4.3.2; R Foundation for Statistical Computing) with the *survival, survminer*, and *ggplot2* packages for Kaplan–Meier and Cox proportional-hazards analyses where complementary to the Python *lifelines* implementation described above. We performed single-cell hierarchical analyses (Methods, “Lifetime analysis of autophagic foci”) in Python 3.11, as detailed in that section. We processed mass spectrometry data in Proteome Discoverer (Thermo Fisher Scientific) using the built-in background-based ANOVA implementation, with downstream filtering and visualization in OriginPro and R. Source data underlying all figures, including individual data points, are provided as Supplementary Data.

## ACKNOWLEDGEMENTS

This work was funded by the NIH NIGMS R35GM156876 to C.B., as well as the Center for Molecular Analysis of Disease Pathways (CMADP) supported by the NIH NIGMS P30GM145499. We thank Luke D. Lavis (Janelia Research Campus, Ashburn, VA, USA) for generously providing the Janelia fluor dyes. We thank Peter McDonald at the Flow Cytometry Core, part of the Chemical Biology of Infectious Disease (CBID) COBRE at the University of Kansas, supported by the National Institute of General Medical Sciences (NIGMS) of the National Institutes of Health under award number P20GM113117. Proteomics data were collected at the Mass Spectrometry and Proteomics Core facility using the Orbitrap Ascend Tribrid System, purchased with funds from the University of Kansas Cancer Center, which is supported by the National Cancer Institute (NCI) Cancer Center Support Grant P30 CA168524.

## AUTHOR CONTRIBUTIONS

T.M.M., S.R.M., and A.G. contributed equally to this work and are listed as first co-authors. C.B. conceived the study, with contributions from D.G.B. C.B. acquired funding and supervised the project. C.B., M.J.R., and A.G. designed the methodology. T.M.M., S.R.M., A.G., U.S., K.N., D.G.B., Z.C., and C.B. performed experiments and acquired data. T.M.M., S.R.M., A.G., Z.C., and C.B. curated data and performed formal analysis. D.G.B. and C.B. developed software. C.B., M.J.R., and S.M.L. provided resources. M.J.R. supervised the proteomics work. T.M.M., S.R.M., A.G., and C.B. validated results. C.B. prepared figures. T.M.M., S.R.M., A.G., and C.B. wrote the original draft. T.M.M., S.R.M., A.G., and C.B. reviewed and edited the final manuscript.

## Supplementary Information

**Fig. S1.**
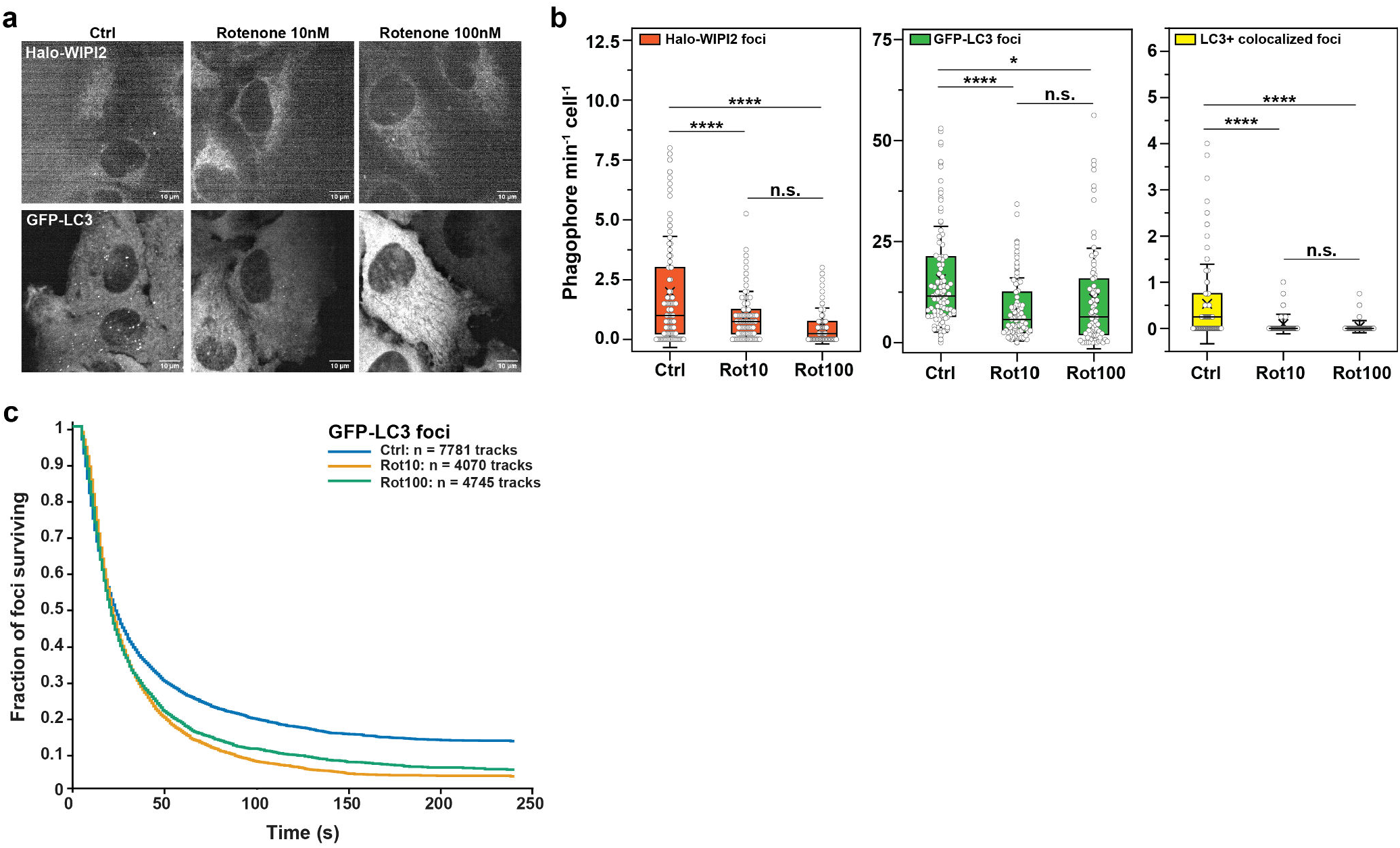
Rotenone suppresses both phagophore initiation and LC3 recruitment. **a**, Representative single frames of live U2OS cells expressing endogenously HaloTagged WIPI2 (top) and GFP-LC3 (bottom), treated for 2 h with vehicle (Ctrl) or rotenone at 10 or 100 nM. Scale bars, 10 µm. **b**, Formation rate of Halo-WIPI2 foci (left), GFP-LC3 foci (centre) and LC3-positive colocalized foci (right), quantified as newly appearing foci per cell per minute. Open circles are individual cells pooled from two independent biological replicates (>100 cells per condition); boxes show the median and interquartile range, whiskers the 10^th^–90^th^ percentiles. Rotenone reduced the formation rate of all three populations at both concentrations, with no further reduction between 10 and 100 nM. P values obtained from ANOVA with Tukey’s post-hoc test (n.s., not significant; *P < 0.05; ****P < 0.0001). **c**, Survival curves for GFP-LC3 foci in the same cells. Curves show the fraction of foci still detected as a function of time after first appearance, pooled across all cells from two independent biological replicates; the number of tracks contributing to each curve is indicated. Rotenone shortened focus lifetime at both concentrations.

**Fig. S2.**
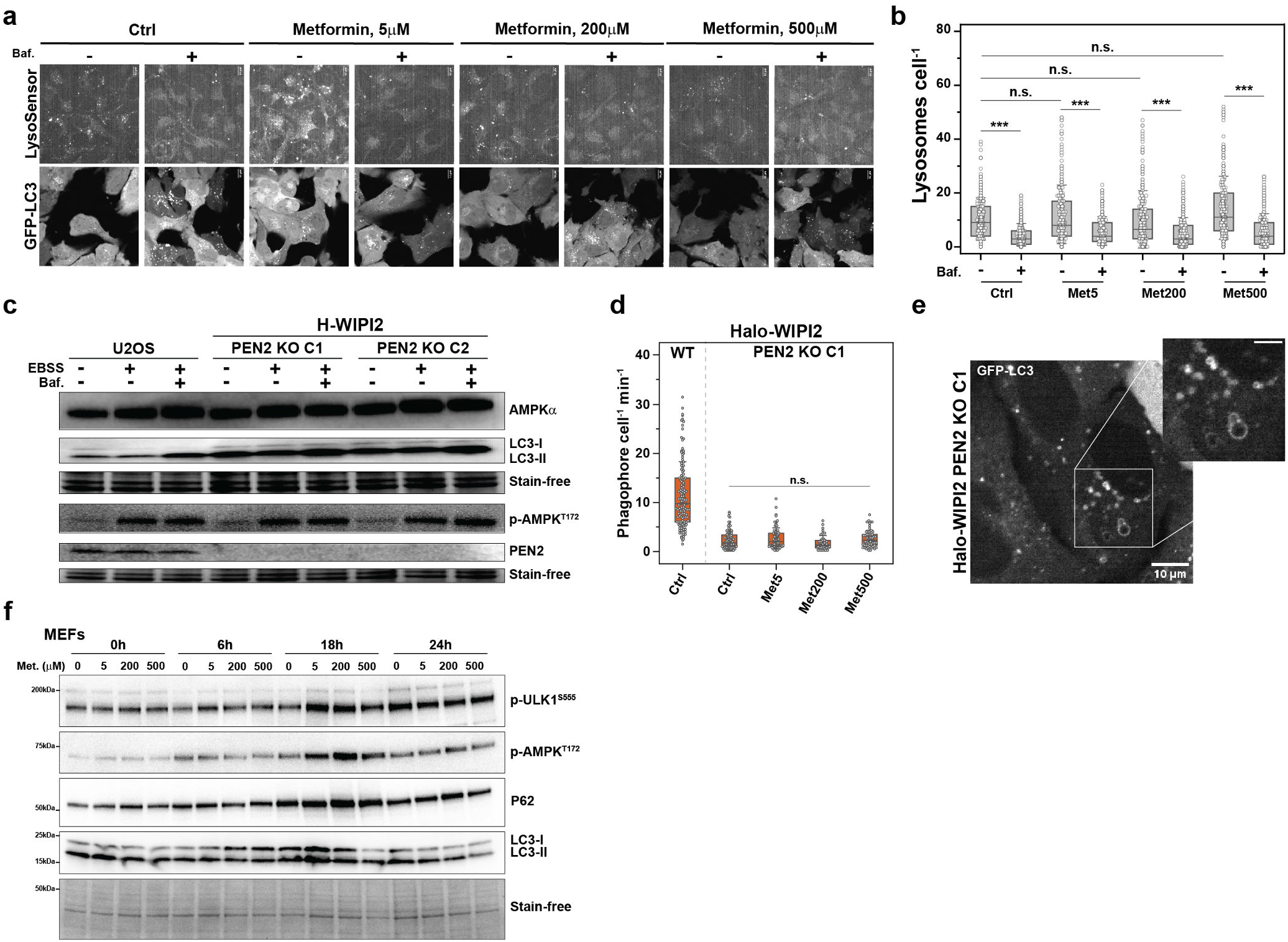
Metformin’s autophagy effects are independent of the lysosomal PEN2 axis. **a**, Representative live-cell images of GFP-LC3 U2OS cells stained with LysoSensor and treated with metformin (5, 200, 500 µM, 2 h) ± bafilomycin A1 (100 nM, 2 h). Scale bar, 10 µm. **b**, Quantification of LysoSensor-positive lysosomes per cell. **c**, Immunoblot of parental U2OS and two independent PEN2 knockout clones (C1, C2) on the Halo-WIPI2 background, treated with EBSS (2 h) ± bafilomycin A1 (100 nM, 1 h). **d**, Halo-WIPI2 phagophore initiation rate (events cell^−1^ min^−1^) in parental H-WIPI2 (Ctrl) and PEN2 KO C1 treated with vehicle or metformin (5, 200, 500 µM, 2 h). **e**, Representative confocal image of GFP-LC3 and Halo-WIPI2 in PEN2 KO C1, showing redistribution of LC3 into large circular structures (inset). Scale bar, 10 µm; inset scale bar, 5 µm. **f**, Immunoblot of mouse embryonic fibroblasts (MEFs) treated with metformin (0, 5, 200, 500 µM) for 0, 6, 18, or 24 h, probed for p-ULK1^S555^, p-AMPK^T172^, P62, and LC3-I/II. Stain-free total protein, loading control. Representative of three independent experiments. Data are shown as box plots (median, 1SD whiskers) with individual cells overlaid. Unpaired two-tailed *t* -test vs. Ctrl, -Baf; n.s., not significant; \*\*\**P* < 0.001. *n* > 25 cells per condition from three independent biological experiments.

**Fig. S3.**
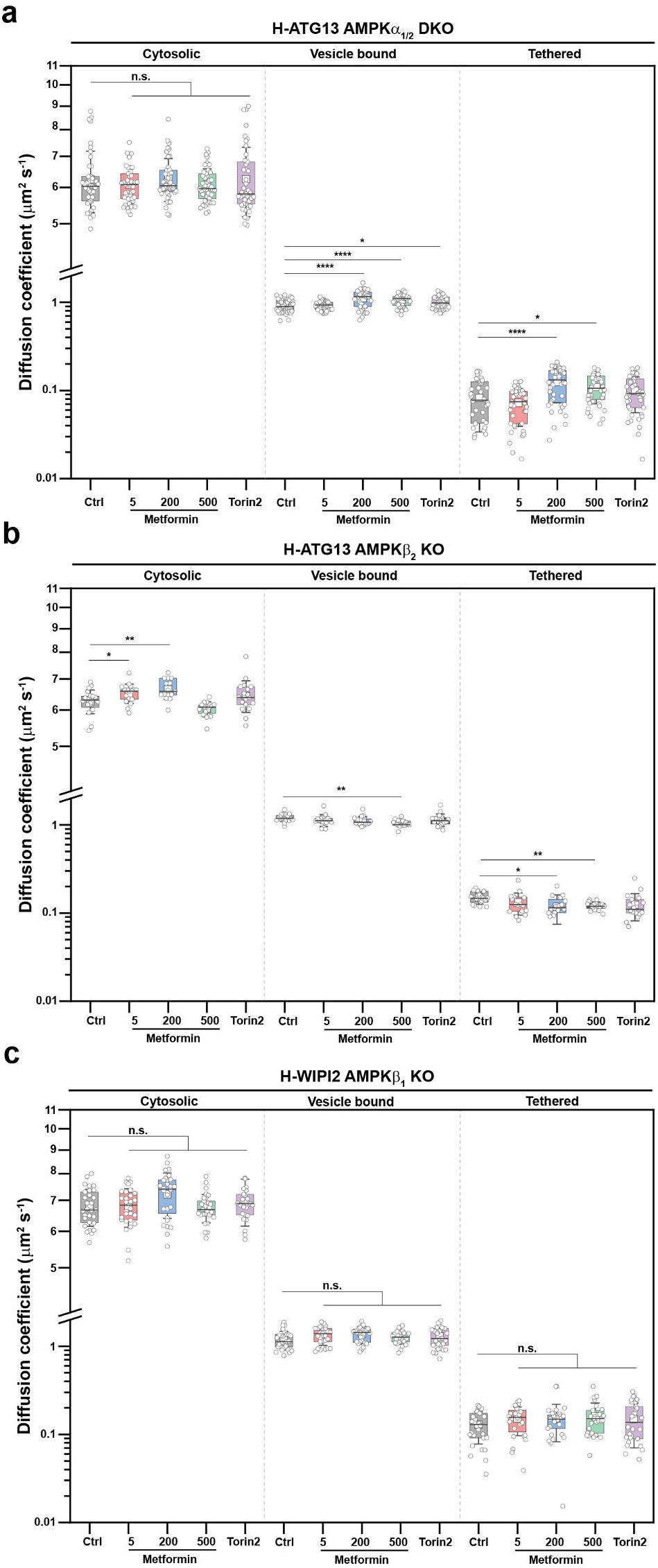
Metformin-induced single-molecule mobility changes in ATG13 and WIPI2 in AMPK-null backgrounds. Single-particle tracking of endogenous Halo-tagged autophagy factors in U2OS cells genetically null for AMPK catalytic or scaffold subunits, treated for 2 h with vehicle (Ctrl), metformin (5, 200, or 500 µM), or Torin2. Trajectories partition into three diffusion states corresponding to a fast cytosolic population (*D* ≈ 5–7 µm^2^ s^−1^), an intermediate vesicle-bound population (*D* ≈ 0.8–1.2 µm^2^ s^−1^), and a slow membrane-tethered population (*D* ≈ 0.05–0.2 µm^2^ s^−1^); diffusion coefficients of individual molecules are plotted on a split log-scale axis. (a) H-ATG13 AMPKα_1/2_ double-knockout cells. **b**, H-ATG13 AMPKβ_2_ knockout cells. **C**, H-WIPI2 AMPKβ_1_ knockout cells. Each circle represents one cell; box plots show median, interquartile range, and Tukey whiskers. Statistical significance was assessed by one-way ANOVA with Dunnett’s post-hoc test against the within-genotype vehicle control. *\*p* < 0.05, *\*\*p* < 0.01, *\*\*\*p* < 0.001, *\*\*\*\*p* < 0.0001; n.s., not significant. n > 15 cells per condition pooled from 3 independent experiments.

**Fig. S4.**
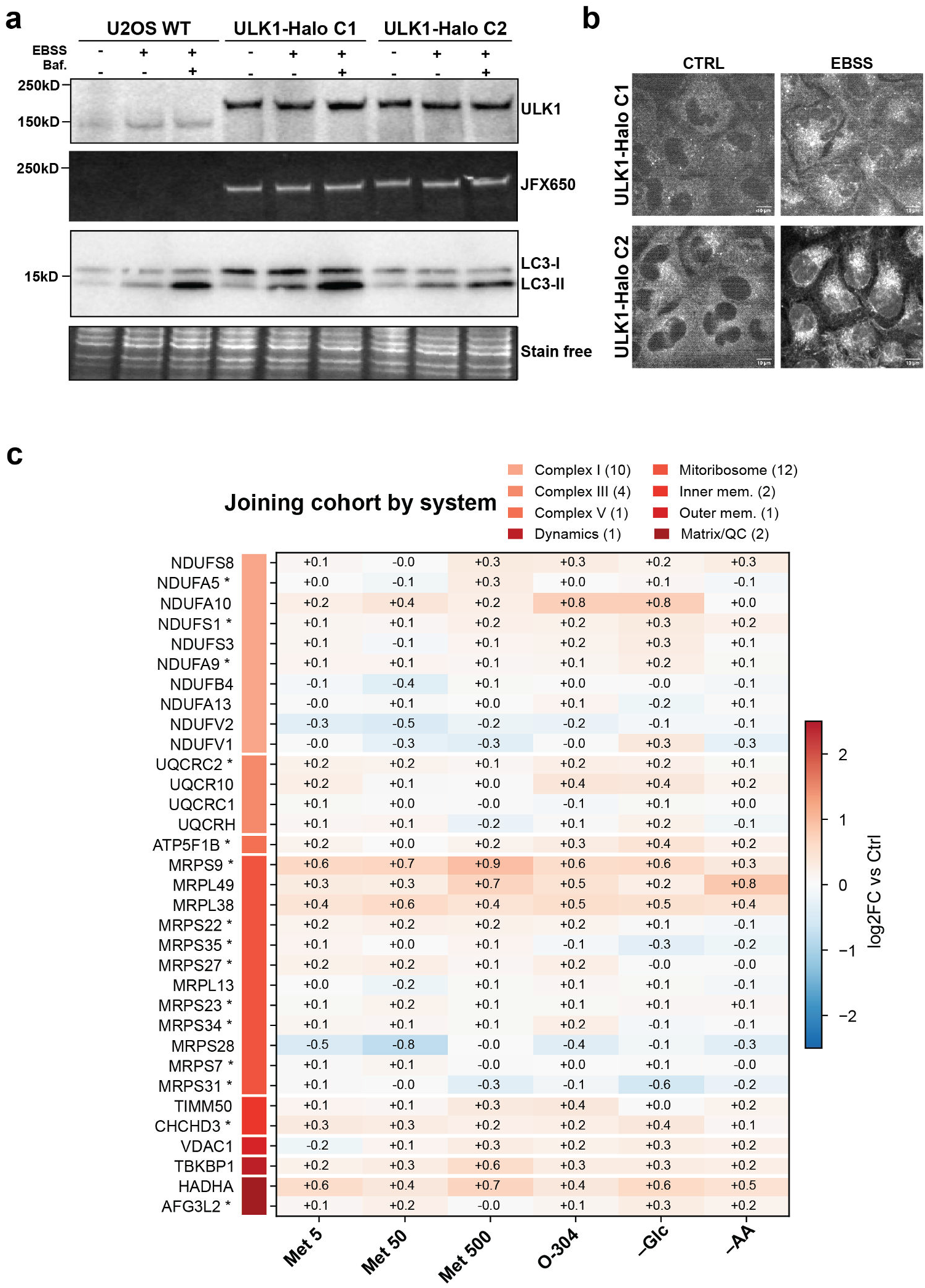
Validation of endogenously Halo-tagged ULK1 in U2OS cells. Two independent CRISPR-edited U2OS clones (ULK1-Halo C1 and ULK1-Halo C2) carrying a homozygous in-frame Halo-tag knock-in at the endogenous *ULK1* locus. **a**, Immunoblot validation. Parental U2OS (WT) and the two knock-in clones were cultured under fed, amino-acid-starved (+EBSS), or starved plus bafilomycin A1 (+EBSS, +Baf.) conditions, then lysed and resolved by SDS-PAGE. The anti-ULK1 blot (top) shows the expected *∼*30 kDa upward shift of the endogenous ULK1 band from *∼*150 kDa in WT to *∼*200 kDa in both knock-in clones, with no detectable signal at the WT molecular weight, confirming biallelic tag insertion. In-gel JFX650 fluorescence (second panel) detects a single band at *∼*200 kDa exclusively in the knock-in lines, demonstrating specific covalent labeling of the tagged protein and absence of free or mistargeted ligand. LC3 immunoblot (third panel) shows EBSS-induced conversion of LC3-I to LC3-II and bafilomycin-trapped LC3-II accumulation in all three lines, confirming that bulk autophagic flux is preserved in the knock-in clones. Stain-free total protein imaging (bottom) is shown as loading control. **b**, Live-cell confocal imaging of ULK1-Halo C1 and C2 cells labeled with 100 nM JFX650 HaloTag ligand, under fed (CTRL) or EBSS conditions. Starvation triggers redistribution of ULK1 from a diffuse cytoplasmic pool into discrete cytoplasmic puncta in both clones, consistent with canonical recruitment of ULK1 to autophagy initiation sites. Scale bars, 10 µm. **c**, Mitochondrial proteins co-purifying with ULK1, grouped by functional system (number of proteins per group in parentheses). Heatmap shows log_2_ fold change versus untreated control for metformin (5, 50, 500 µM), O-304, glucose withdrawal (-Glc) and amino-acid withdrawal (-AA); n = 3 biological replicates. Asterisks indicate significance *P* > 0.05.

### Supplementary Movie Legends

**Movie S1**. Representative live-cell movie of U2OS cells endogenously expressing 3×FLAG-HaloTag-ATG13 (JFX650) and GFP-LC3, treated with vehicle, bafilomycin A1 (Baf), O-304, or O-304 + Baf. Acquired at 1 frame s^−1^; 650 × 650 pixels, 0.217 µm/pixel.

**Movie S2**. Representative live-cell movie of U2OS cells endogenously expressing 3×FLAG-HaloTag-ATG13 (JFX650) and GFP-LC3, treated with metformin (5 µM), metformin + bafilomycin A1 (Baf), metformin + O-304, or metformin + O-304 + Baf. Acquired at 1 frame s^−1^; 650 × 650 pixels, 0.217 µm/pixel.

**Movie S3**. Representative live-cell movie of U2OS cells endogenously expressing 3×FLAG-HaloTag-ATG13 (JFX650) and GFP-LC3, treated with metformin (200 µM), metformin + Baf, metformin + O-304, or metformin + O-304 + Baf. Acquired at 1 frame s^−1^; 650 × 650 pixels, 0.217 µm/pixel.

**Movie S4**. Representative live-cell movie of U2OS cells endogenously expressing 3×FLAG-HaloTag-ATG13 (JFX650) and GFP-LC3, treated with metformin (500 µM), metformin + Baf, metformin + O-304, or metformin + O-304 + Baf. Acquired at 1 frame s^−1^; 650 × 650 pixels, 0.217 µm/pixel.

**Movie S5**. Representative live-cell movie of U2OS cells endogenously expressing 3×FLAG-HaloTag-WIPI2 (JFX650) and GFP-LC3, treated with vehicle, bafilomycin A1 (Baf), O-304, or O-304 + Baf. Acquired at 1 frame s^−1^; 650 × 650 pixels, 0.217 µm/pixel.

**Movie S6**. Representative live-cell movie of U2OS cells endogenously expressing 3×FLAG-HaloTag-WIPI2 (JFX650) and GFP-LC3, treated with metformin (5 µM), metformin + bafilomycin A1 (Baf), metformin + O-304, or metformin + O-304 + Baf. Acquired at 1 frame s^−1^; 650 × 650 pixels, 0.217 µm/pixel.

**Movie S7**. Representative live-cell movie of U2OS cells endogenously expressing 3×FLAG-HaloTag-WIPI2 (JFX650) and GFP-LC3, treated with metformin (200 µM), metformin + Baf, metformin + O-304, or metformin + O-304 + Baf. Acquired at 1 frame s^−1^; 650 × 650 pixels, 0.217 µm/pixel.

**Movie S8**. Representative live-cell movie of U2OS cells endogenously expressing 3×FLAG-HaloTag-WIPI2 (JFX650) and GFP-LC3, treated with metformin (500 µM), metformin + Baf, metformin + O-304, or metformin + O-304 + Baf. Acquired at 1 frame s^−1^; 650 × 650 pixels, 0.217 µm/pixel.

**Movie S9**. Representative live-cell movie of U2OS cells expressing GFP-LC3, untreated or treated with metformin (5, 200 and 500 µM) with and/or without DMS (1 mM). Acquired at 1 frame s^−1^; 350 × 350 pixels, 0.217 µm/pixel.

**Movie S10**. Representative live-cell movie of U2OS cells expressing GFP-LC3, untreated or treated with metformin (5, 200 and 500 µM) with and/or without α-KB (0.5 mM). Acquired at 1 frame s^−1^; 350 × 350 pixels, 0.217 µm/pixel.

**Movie S11**. Representative live-cell movie of U2OS cells endogenously expressing 3×FLAG-HaloTag-ATG13 (JFX650) and GFP-LC3 and having AMPK α_1/2_ double knock-out, untreated or treated with metformin (5 and 50 µM) with and/or without O-304 (1 µM). Acquired at 1 frame s^−1^; 350 × 350 pixels, 0.217 µm/pixel.

**Movie S12**. Representative live-cell movie of U2OS cells endogenously expressing 3×FLAG-HaloTag-ATG13 (JFX650) and GFP-LC3 and having AMPK β_2_ knock-out, untreated or treated with metformin (5, 200 and 500 µM) with and/or without O-304 (1 µM). Acquired at 1 frame s^−1^; 350 × 350 pixels, 0.217 µm/pixel.

**Movie S13**. Representative live-cell movie of U2OS cells endogenously expressing 3×FLAG-HaloTag-WIPI2 (JFX650) and GFP-LC3 auntreated or treated with rotenone (10 and 100 nM) with and/or without O-304 (1 µM). Acquired at 1 frame s^−1^; 350 × 350 pixels, 0.217 µm/pixel.

